# Multiple HP1 binding motifs in SUV420H2 heterochromatic targeting module

**DOI:** 10.1101/2023.12.03.569804

**Authors:** Masaru Nakao, Yuko Sato, Arisa Aizawa, Hiroshi Kimura

## Abstract

Constitutive heterochromatin is transcriptionally repressed and densely packed chromatin, typically harboring histone H3 Lys9 trimethylation (H3K9me3) and heterochromatin protein 1 (HP1). SUV420H2, a histone H4 Lys20 methyltransferase, is recruited to heterochromatin by binding to HP1 through its Heterochromatic Targeting Module (HTM). Here, we have identified three HP1 binding motifs within the HTM. Both the full-length HTM and its N-terminal region (HTM-N), which contains the first and second motifs, stabilized HP1 on heterochromatin. The intervening region between the first and second HP1 binding motifs in HTM-N was also crucial for HP1 binding. In contrast, the C-terminal region of HTM (HTM-C), containing the third motif, destabilized HP1 on chromatin. An HTM V374D mutant, featuring a Val374 to Asp substitution in the second HP1 binding motif, localizes to heterochromatin without affecting HP1 stability. These data suggest that the second HP1 binding motif in the SUV420H2 HTM is critical for locking HP1 on H3K9me3-enriched heterochromatin. HTM V374D, tagged with a fluorescent protein, can serve as a live-cell probe to visualize HP1-bound heterochromatin.

## Introduction

Heterochromatin is a form of densely packed chromatin in which transcription is typically suppressed. Constitutive heterochromatin harboring histone H3 Lys9 trimethylation (H3K9me3) is often formed on DNA repeat elements. Heterochromatin protein 1 (HP1) binds to both H3K9me2 and H3K9me3 through its chromodomain (CD) (Bannister et al., 2001; Lachner et al., 2001; Nielsen et al., 2002). In mammalian cells, there are three subtypes of HP1, namely HP1α, HP1β, and HP1γ (Jones et al., 2000). HP1α and HP1β are predominantly found in pericentromeric heterochromatin, also known as chromocenters, in mouse cells, whereas HP1γ is generally localized in euchromatin (Nielsen et al., 2001; Minc et al., 2000). The dimerization of HP1 through its chromoshadow domain (CSD) is implicated in chromatin compaction, as it facilitates the bridging of two nucleosomes within H3K9me3-rich chromatin fibers (Hiragami-Hamada et al., 2016; Machida et al., 2018) and provides a binding platform for effector proteins that contain PxVxL motif (Smothers & Henikoff, 2000; Nozawa et al., 2010; Yan et al., 2018). Furthermore, HP1 is also thought to contribute to heterochromatin compaction through phase separation (Strom et al., 2017; Larson et al., 2017; Sanulli et al., 2019; Qin et al., 2021).

H4K20me3 is also enriched in constitutive heterochromatin, in a manner dependent on H3K9me3 and HP1 (Schotta et al., 2004; Hahn et al., 2013). Four histone methyltransferases – SUV420H1, SUV420H2, SMYD5, and SMYD3 – have been reported to mediate the H4K20me3 modification (Schotta et al., 2004; Kidder et al., 2017; Stender et al., 2012; Foreman et al., 2011). Among these, SUV420H2 is thought to be the primary enzyme responsible for H4K20me3, as SUV420H2 knock out (KO) results in an almost complete loss of this modification (Schotta et al., 2008). Although Suv4-20h2 KO mice develop normally (Schotta et al., 2008), cells lacking SUV420H2 show reduced heterochromatin compaction, chromocenter fragmentation, and abnormalities in mitotic spindle separation during mitosis (Hahn et al., 2013). The minimal effect of Suv4-20h2 KO in development could be due to its low expression during the embryonic development stage and/or a potential compensatory effect by Suv4-20h1, which possesses a SET domain highly similar to that of SUV420H2 (Schotta et al., 2008). In vitro studies have indeed demonstrated that the SET domain of SUV420H1 is capable of mediating both H4K20me2 and me3 modifications (Schotta et al., 2004). While the loss of SUV420H2 does not significantly affect mouse development, ectopic expression of SUV420H2 in pre-implantation embryos disrupts embryogenesis, indicating defects in the S-phase progression (Eid et al., 2016). Artificial enrichment of H4K20me3 at centromeres restricts the localization and function of AuroraB, leading to an increase in errors during chromosome separation in mitosis (Herlihy et al., 2021). Hence, controlled H4K20me3 localization appears to be important in regulating the genome function.

The mechanism by which H4K20me3 and SUV420H2 localize to pericentromeric heterochromatin, or chromocenters, in mouse cells has been investigated. SUV420H2 contains a region known as the Heterochromatic Targeting Module (HTM) at its C-terminal region, which binds to HP1 on H3K9me3-enriched chromatin (Schotta et al., 2004; Hahn et al., 2013). Interestingly, a photobleaching assay indicated that HTM tagged with the green fluorescent protein (GFP) binds to chromatin more stably than HP1-GFP (Souza et al., 2009; Hahn et al., 2013). Such different behavior can be attributed to the multiple HP1 binding modules in the HTM (Souza et al., 2009). When HTM is divided into two fragments, each independently binds to HP1 and accumulates at chromocenters (Hahn et al., 2013). However, the detailed mechanism of how the SUV420H2 HTM binds to HP1 remains elusive.

In this study, we analyzed the localization and HP1 binding of various deletion and amino acid substitution mutants of the SUV420H2 HTM. We identified the three HP1 binding motifs whose spatial configuration was crucial for efficient heterochromatin targeting and HP1 interaction when overexpressed. One of the HTM mutant which harbors two independent HP1 binding motifs could serve as a live-cell probe for detecting HP1 on heterochromatin.

## Results

### SUV420H2 harbors three HP1-binding motifs within its heterochromatic targeting module (HTM)

Previous studies have indicated that two regions within the HTM of SUV420H2 bind to HP1 (Hahn et al., 2013), yet the specific binding motifs have not been defined. To identify the HP1 binding regions within the HTM, we constructed various truncated mutants and transiently expressed them in mouse A9 cells (Fig. S1). The full HTM tagged with superfolder GFP (HTM-sfGFP) localized to pericentromeric heterochromatin, or chromocenters, highlighted by H3K9me3 and Hoechst 33342 staining (Fig. S1A and S1B), as reported previously (Hahn et al., 2013). In cells expressing HaloTag-tagged KDM4D, an H3K9me2/3 demethylase (Hayashi-Takanaka et al., 2020), HTM-sfGFP was diffused throughout the nucleoplasm (Fig. S1C), indicating that the heterochromatin targeting of HTM is dependent on H3K9me2/3 to which HP1 binds. Both the N- and C-terminal regions of HTM, amino acid (aa) 347-380 and 381-435, respectively, could target heterochromatin (Hahn M 2013), albeit more weakly than the full HTM (Fig. S1D). We thus constructed deletion mutants to determine the minimal essential element required for chromocenter enrichment in each N- or C-region (Fig. S1D). Within the HTM N-terminal region, aa 352-380 exhibited chromocenter localization (heterochromatin enrichment ratio 2.59 vs. 2.67 for aa 347-380), whereas a further N-terminal deletion (aa 356-380) distributed homogenously in the nucleoplasm, and a further C-terminal deletion (aa 352-374) was poorly enriched at chromocenters (heterochromatin enrichment ratio 1.65) (Fig. S1D). Regarding to the C-terminal region, aa 389-410 still localized to chromocenters (heterochromatin enrichment ratio 1.73 vs 1.87 for aa 381-435). A further C-terminal deletion mutant (aa 381-404) distributed homogenously throughout the nucleoplasm, and a further N-terminal deletion mutant (391-410) showed reduced accumulation at chromocenters (heterochromatin enrichment ratio 1.39). Since aa 352-380 and 389-410 retained the similar chromocenters targeting efficiency to their respective original fragments, we selected these for further point mutation studies and designated them as the HTM-N and HTM-C, respectively.

A previous study has indicated that the full HTM (aa 347-435) binds to HP1 proteins through CSD (Souza et al., 2009), but the exact amino acids that are required for HP1 binding were not defined. To identify HP1 binding motifs within the HTM-N and HTM-C regions, we aligned amino acid sequences from different species, as essential amino acids are often evolutionally covered (Fig. S2). In the HTM-N region, two PxVxL-like motifs were found at aa 352-356 (ARVSL in humans) and 372-376 (ALVAL in humans). These motifs were moderately conserved across amniota, including *Homo sapiens*, *Mus musculus*, *Phascolarctos cinereus*, *Ornithorhynchus anatinus*, *Alligator mississippiensis*, *Chrysemys picta*, *Anolis carolinensis*, and *Gallus gallus* (Fig. S2A and S2B). In the HTM-C region, a motif combining PxVxL- and PxxVxL-like sequences (Yanli et al., 2017, Maeda et al., 2022) was found at aa 389-408 (PYVVRVDL in humans), which was conserved only among mammals (Fig. S2A and S2B). To confirm that these HP1 binding motifs are necessary for chromocenter targeting, we engineered amino acid substitution mutants by replacing Val to Asp (Fig. 1) because it has been reported that a V to D substitution in the PxVxL disrupt the hydrophobic interaction with HP1 (Vassallo et al., 2002, Yanli et al., 2017). The V354D mutant of the HTM-N distributed homogenously throughout the cells (Fig. 1A and 1B), indicating that this mutation abolished the interaction between the HTM-N and HP1. Conversely, the V374D mutant still accumulated in the chromocenters, albeit the enrichment ratio was substantially lower compared to the wild-type HTM-N (1.59 vs 2.59) (Fig. 1B and 1C). This suggests that the PxVxL-like motif at aa 352-356 (ARVSL) alone binds to HP1 and the motif at aa 372-376 (ALVAL) assist in a more stable binding. Regarding the HTM-C, mutations V402D, V404D, and the double V402D-V404D mutations eliminated chromocenter localization (Fig. 1D), demonstrating that both Val residues within the HTM-C are crucial for HP1 interaction. A full HTM mutant containing the triple mutation V354D-V402D-V404D did not accumulate at chromocenters (Fig. 1E), indicating that there is no additional HP1 binding motifs that could mediate heterochromatin targeting within SUV420H2 HTM. Moreover, the heterochromatin enrichment ratios of the HTM mutants harboring single V354D and the double V402D-V404D mutations (2.05 and 2.95, respectively) were comparable to those of the HTM-C (1.73) and HTM-N (2.59), respectively (Fig, 1C and 1E), supporting the view that the SUV420H2 HTM binds to HP1 through three HP1 binding motifs.

**Fig. 1.**
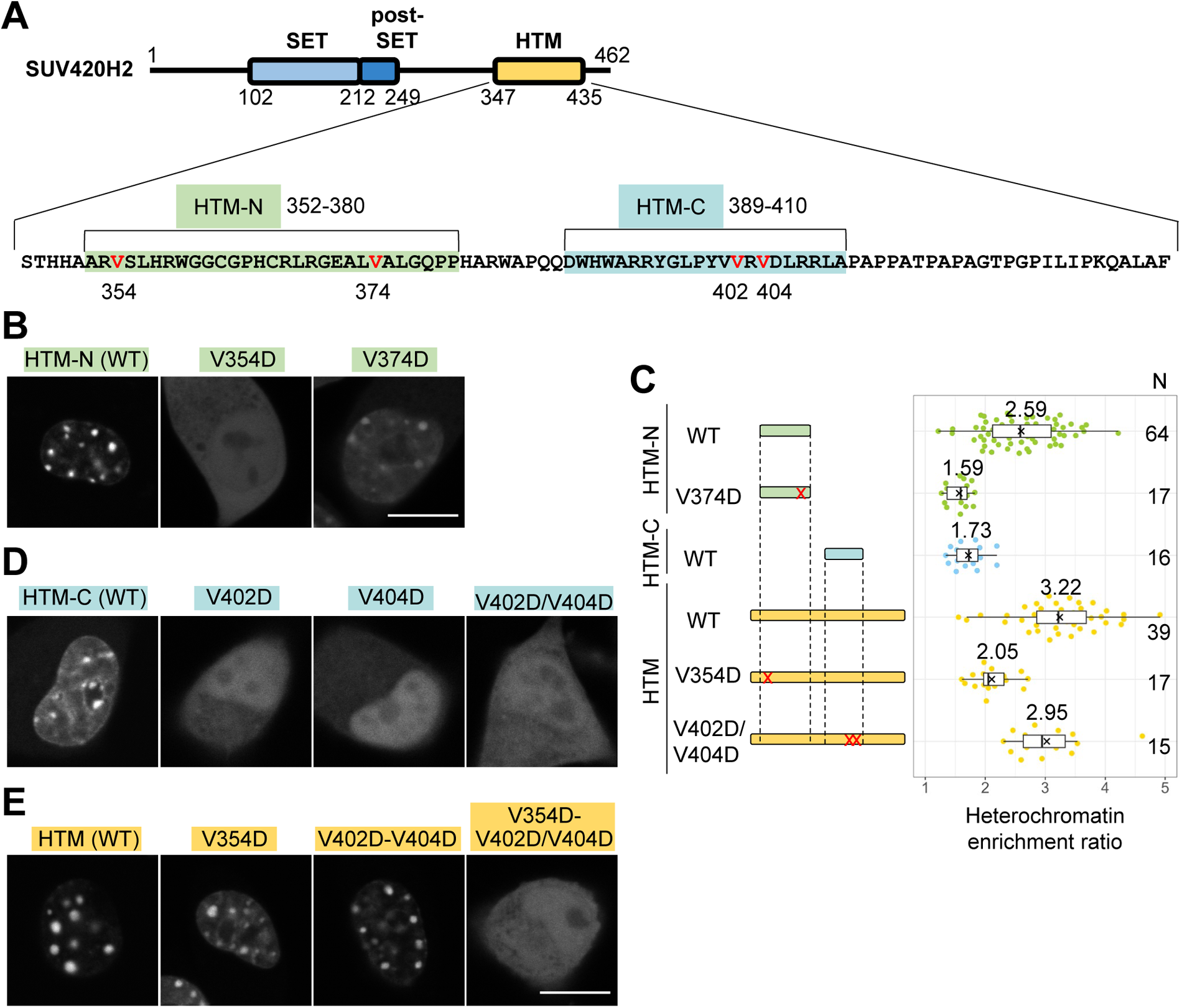
Human SUV420H2 contains three HP1 binding sites within HTM. (A) Schematic representation of SUV420H2 domains, including the amino acid sequence of HTM. The N- and C-terminal heterochromatic targeting regions (aa 352-380 and 389-410) are labeled as HTM-N and HTM-C, respectively. Val residues in HP1 binding motifs are indicated in red. (B-E) Analysis of the impact of Val to Asp mutations in HP1-binding PxVxL-like motifs on heterochromatin localization. sfGFP-tagged HTM-N (B), HTM-C (D), and full HTM (E), along with their mutants, were expressed in A9 cells. Heterochromatin accumulation of each sfGFP-tagged protein is represented as heterochromatin enrichment ratio relative to the nucleus (C). (B, D, and E) Single confocal sections of typical nuclei. (C) Boxplots. Center lines represent the medians; box limits indicate the 25th and 75th percentiles; whiskers extend 1.5 times the interquartile range from the 25th to 75th percentiles; × denote the means; and individual data points are shown as dots. Median values and number of analyzed nuclei are indicated above and to the right of box plots, respectively, respectively. Scale bars: 10 μm.

To investigate the interaction between HP1 and the HTM-N and HTM-C regions, as well as the impact of mutations within HP1-binding motifs, we carried out co-immunoprecipitation assays using cell extracts prepared from HeLa cells that express sfGFP, HTM-N-sfGFP, HTM-C-sfGFP, and their corresponding Val to Asp mutants, namely HTM-N V354D and HTM-C V402D-V404D (Fig. 2A and 2B). During preparation of cell lysates using a high salt buffer containing 1 M NaCl, HP1 proteins, especially HP1β, were less efficiently extracted from cells expressing HTM-N-sfGFP, along with the HTM-N-sfGFP itself (Fig. 2A, lanes 5 and 14), compared to those expressing sfGFP and HTM-N V354D-sfGFP (Fig. 2A, lanes 4 and 13, and 6 and 15). This suggests that the expression of HTM-N may stabilize the association of HP1 with chromatin (a point that will be discussed later). Following immunoprecipitation using the cell extracts with anti-GFP magnetic beads, the recovery of the sfGFP fusion proteins and HP1 subtypes was assessed by immunoblotting. All three HP1 subtypes, HP1α, HP1β, and HP1γ, were co-immunoprecipitated with both HTM-N-sfGFP (Fig. 2A, lane 11) and HTM-C-sfGFP (Fig. 2B, lane 8), but not with their corresponding mutants (V354D and V402-V404D) nor with sfGFP alone (Fig. 2A, lane 10 and 12, and 2B, lane 7 and 9). Despite the HTM-N V354D mutant retaining a PxVxL-like motif at aa 372-376, this motif on its own does not appear to suffice to form a stable complex with HP1. These results, aligning with microscopic data to evaluate heterochromatin targeting, reinforced the conclusion that the SUV420H2 HTM is directed to chromocenters via its interaction with HP1 CSD (Souza et al., 2009).

**Fig. 2.**
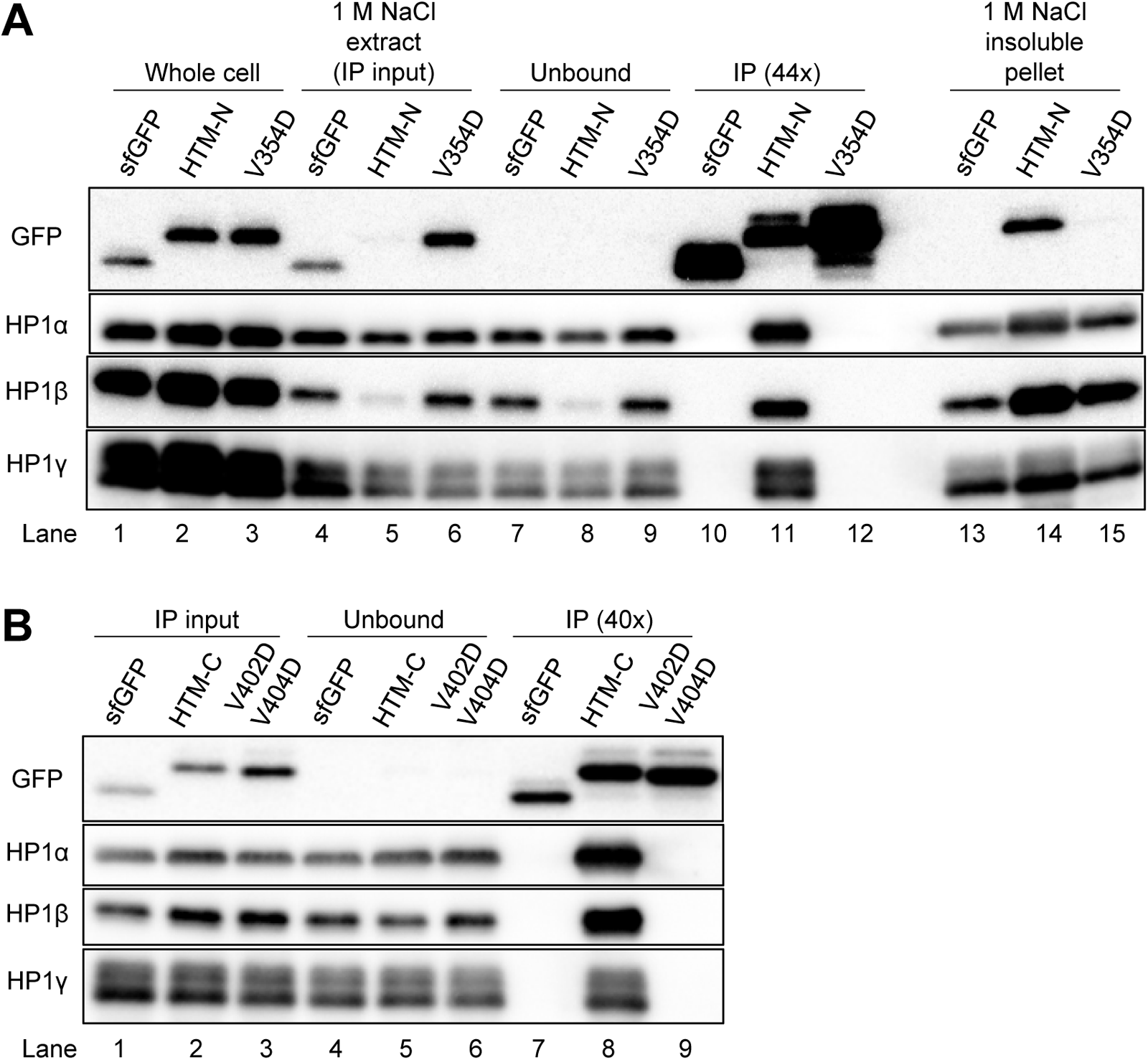
HTM-N and HTM-C interaction with HP1 through their PxVxL-like motifs. (A) Analysis of HeLa cells expressing sfGFP, HTM-N-sfGFP, and HTM-N V354D-sfGFP (in which Val354 is substituted with Asp). Cells were treated with a buffer containing 0.1% Triton X-100 and 1 M NaCl to extract most chromatin-binding proteins (whole cell fractions; lanes 1-3). Post-centrifugation, the supernatants (1 M NaCl extract; lanes 4-6) were separated from the pellet (1 M NaCl pellet; lanes 13-15) and utilized as inputs for immunoprecipitation using anti-GFP beads. The unbound fractions (Unbound; lanes 7-9) and the immunoprecipitated materials (IP; 40x concentrated compared to the input; lanes 10-12) were processed. Protein samples were then separated on SDS-polyacrylamide gels, transferred to membranes, and probed with the indicated antibodies. (B) Extracts from HeLa cells expressing sfGFP, HTM-C-sfGFP, and HTM-C V402D-V404D-sfGFP were prepared, immunoprecipitated, and immunoblotted, as described in (A). The full blots are shown in Supplementary Fig. S8.

### Zinc finger-like motif in HTM-N is crucial for targeting to heterochromatin

In HTM-N, three amino acids, H357, C362, and C366, positioned between the two PxVxL-like motifs, were highly conserved across different species (Fig. S2B). To examine whether these conserved aa residues are essential for heterochromatin targeting, we expressed HTM-N-sfGFP H357A, C362V, and C366V single mutants in A9 cells (Fig. 3A). Each of the three mutants distributed throughout the nucleoplasm and cytoplasm (Fig. 3A), implying that each of these His and Cys residues is crucial for HP1 binding. Given the importance of His and Cys, we postulated that this region might adopt a structure akin to a zinc finger domain with four Cys/His residues (Neuhaus et al., 2022) to properly orient the two PxVxL-like motifs, thereby bridging two HP1 CSD binding sites and stabilizing the interaction. To evaluate this hypothesis, we constructed a mutant that included an insertion of a flexible linker comprising a 3x(GGGGS) sequence and then expressed it in A9 cells (Fig. 3B). Although this mutant localized to chromocenters, its enrichment was lower compared to HTM-N, similarly to the mutant lacking the second PXVXL-like motif (Fig. 3B). Thus, in addition to the critical functions of the His and Cys residues in HP1 binding, a specific configuration of the intervening region appears to assist in cooperative binding of the two HP1 binding motifs to stabilize the interaction.

**Fig. 3.**
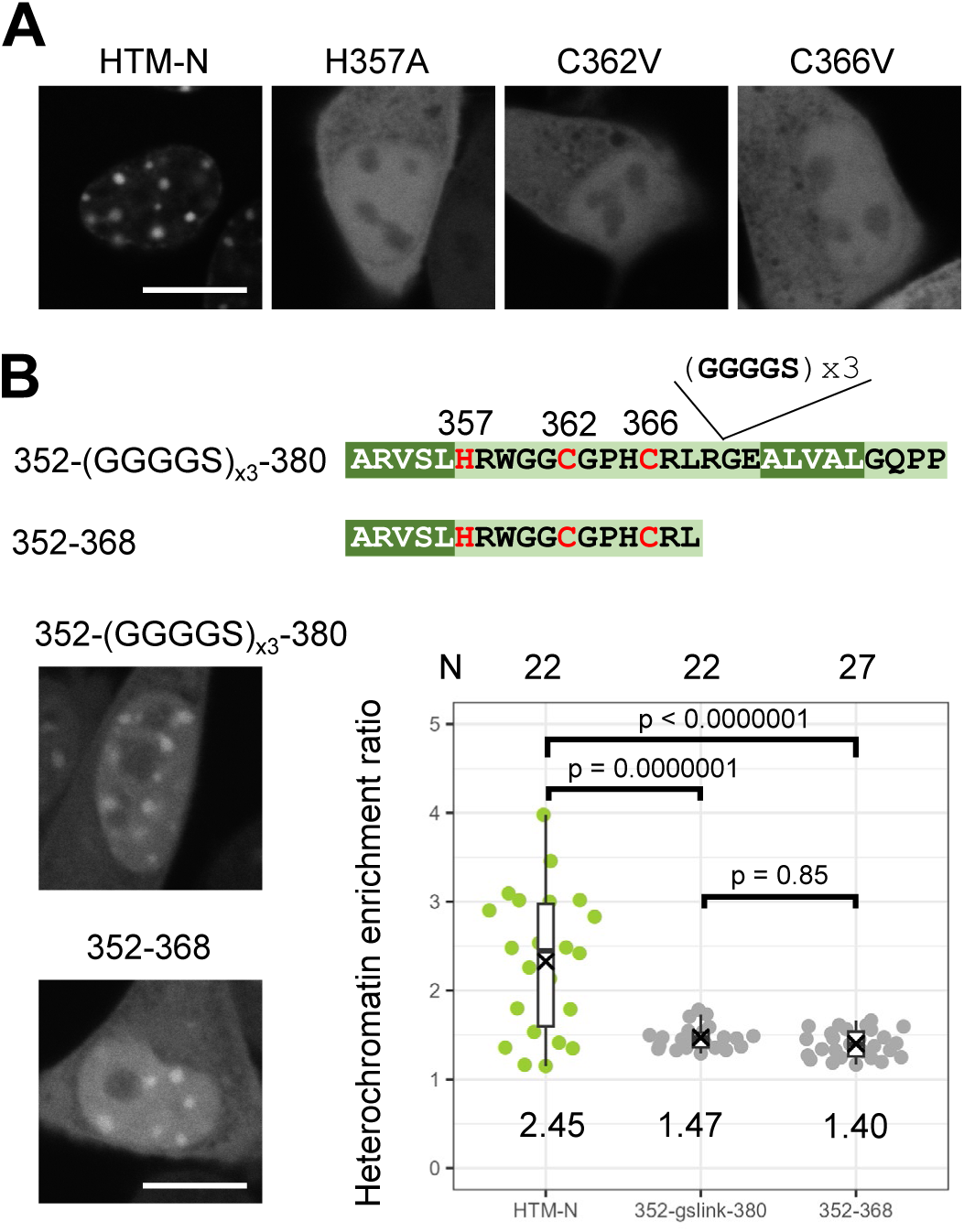
Functions of the linker region between two HP1 binding motifs in HTM-N. (A) The evolutionally conserved His and Cys resides are critical for HTM-N heterochromatin targeting. Single amino acid substitution mutants of HTM-N (H357A, C362V, and C368V) were expressed in A9 cells and their localization was analyzed using confocal microscopy. (B) The heterochromatic accumulation of HTM-N is weakened by the insertion of a flexible linker between the two HP1 binding motifs and the deletion of the second motif. Single confocal sections of typical nuclei are shown, along with quantitative data as in Fig. 1. The averages of enrichment ratios and the numbers of analyzed nuclei are indicated above and below box plots, respectively. p-values by Tukey-Kamer test are indicated. Scale bars: 10 μm.

### Overexpression of HTM-sfGFP facilitates HP1β accumulation in H3K9me3-enriched heterochromatin

HP1 at chromocenters can be maintained and immobilized by “HP1 locking” with two HP1 binding modules within a protein, like SENP7 (Romeo et al., 2015). SUV420H2 HTM-N was implied to cause HP1 locking because HP1 exhibited resistant to extraction, as demonstrated by immunoblotting (Fig. 2A). To investigate the impact of HTM expression on HP1 at the cellular level, HP1β signals in wild-type and HTM-sfGFP-expressing cells were compared. For this purpose, we used human HeLa cells because HP1 is highly enriched in chromocenters in mouse cells and a further enrichment by HP1 locking may not be easily detected. To directly compare cells without and with HTM-sfGFP expression in the same microscopic field, HeLa cells stably expressing HTM-sfGFP were co-plated with the parental cells and stained with antibodies specific to HP1β and H3K9me3. HP1β was more pronouncedly concentrated in cells expressing HTM-sfGFP compared to wild-type cells (Fig. 4A, 4B, and S3), without affecting H3K9me3 levels (Fig. S3B and S3C). Quantitative measurements of the enrichment ratios of HP1β signals in H3K9me3-enriched heterochromatin (in the top 10% highest H3K9me3 intensity pixels over the whole nucleus) revealed that the enrichment ratio was substantially higher in cells expressing HTM-sfGFP. This result is consistent with the immunoblotting data showing increased levels of the total and extraction-resistant HP1β in cells expressing HTM-sfGFP (Fig. 2A). A weaker HP1-locking effect was observed when HTM-N, which has two PxVxL-like motifs, was expressed (Fig. 4A and 4B). Conversely, the expression of HTM-C slightly decreased HP1 accumulation (Fig. 4A and 4B). Note that the HTM-C and HTM V374D, which is described below, were diffusely distributed throughout the nucleus in paraformaldehyde-fixed cells, likely because their binding to HP1 is not very stable and cross-linkable Lys residues are limited. We also examined the effect of an HTM mutant that harbors a Val to Asp substitution in the second PxVxL-like motif, necessary for stable HP1 binding in conjunction with the first PxVxL-like motif and the zinc-finger-like motif. Although this mutant, HTM V374D, retained two HP1 binding motifs (aa 352-356 and 399-408), no HP1 locking effect was observed (Fig. 4A and B). These results thus suggest that the second HP1 binding motif at 372-376 (ALVAL) plays a role in HP1 locking and merely having two HP1 binding sites is insufficient for HP1 locking. Thus, proper configuration of multiple binding modules appears to be crucial in HP1 locking.

**Fig. 4.**
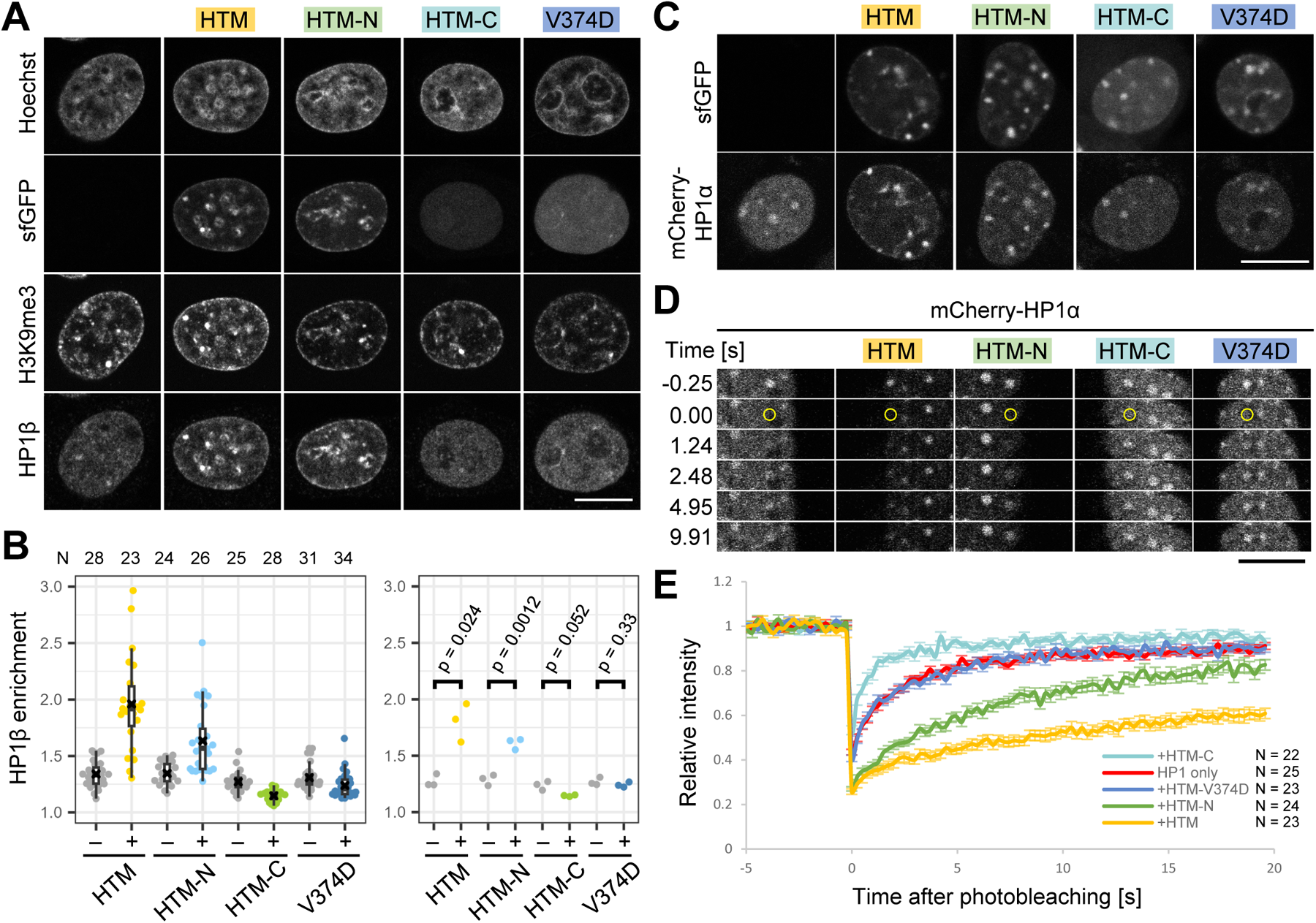
Expression of HTM-sfGFP and HTM-N-sfGFP facilitates HP1β accumulation in H3K9me3-enriched heterochromatin. (A and B) Effects of SUV420 HTM expression on HP1β accumulation in H3K9me3-enriched heterochromatin. HeLa cells expressing HTM, HTM-N, HTM-C, and HTM V374D, each tagged with sfGFP, were stained with specific antibodies directed against H3K9me3 and HP1β. DNA was counterstained with Hoechst 33342. (A) Single confocal sections of typical nuclei. See Fig. S4 for larger fields of view containing multiple nuclei. (B) Quantification of HP1β on H3K9me3-marked heterochromatin. Using images like those shown in (A), the average HP1β intensities in the top 10% of H3K9me3 highest intensity pixels were measured and normalized to the mean intensities in entire nuclei. HP1β enrichments in H3K9me3-enriched heterochromatin in individual nuclei from a single experiment are shown as dot and box plots on the left. The numbers of nuclei (*N*) analyzed are indicated above the box plots. Mean values of individual experiments in biological triplicates are shown on the right. Between 17-37 nuclei were analyzed for each mutant in single experiments. p-values obtained from Student’s t-test (paired, two-tailed) are indicated. (C and D) Co-expression of sfCherry-HP1α with HTM, HTM-N, HTM-C, and HTM V374D, each tagged with sfGFP. (C) Representative images of sfCherry-HP1α in live cells when co-expressed with HTM and its mutants are displayed. (D) Fluorescence recovery after photobleaching. During time-lapse imaging, an area containing sfCherry-HP1α focus (yellow circle) was bleached (top panels). The graphs below show the relative fluorescence intensities of sfCherry-HP1α in the bleached area, normalized by those before bleaching (mean ± s.e.m.). The total number of cells (*N*) analyzed from two independent experiments are shown. Scale bars: 10 μm.

We next investigated whether the effect on HP1β induced by overexpression of HTM-N-sfGFP is also observed for HP1α and HP1γ. HeLa cells stably expressing HTM-N-sfGFP were co-plated with and the parental cells and then stained with antibodies against HP1α, HP1β, and HP1γ. The intensity of each HP1 subtype was higher in cells expressing HTM-N-sfGFP compared to the control parental cells while HP1β showed the highest intensity increase (Fig. S4), which is in good agreement with the immunoblotting data (Fig. 2).

Finally, we analyzed the effects of HP1 locking using fluorescence recovery after photobleaching (FRAP) (Romeo et al., 2015; Souza et al., 2009; Hahn et al., 2013) using sfCherry-HP1α. When sfCherry-HP1α was co-expressed with the full HTM in HeLa cells, sfCherry-HP1α appeared to be more concentrated at heterochromatin compared to cells without the full HTM (Fig. 4C) and the fluorescence recovery rate was drastically decreased (Fig. 4D). The recovery of sfCherry-HP1α was also slowed when co-expressed with HTM-N-sfGFP, albeit the effect was less pronounced than with the full HTM-sfGFP (Fig. 4D). In contrast, when HTM-C-sfGFP was expressed, the recovery of sfCherry-HP1α was slightly accelerated (Fig. 4D), which aligns with the effects of the expression of single PxVxL module in SENP7 (Romeo et al., 2015). The recovery rate of sfCherry-HP1α was not affected by HTM V374D-sfGFP expression. These data are in complete agreement with the immunofluorescence results described above.

### Depletion of SUV420H1 and SUV420H2 did not affect HP1 signals

To investigate whether endogenous SUV420H2 affects HP1 locking, as observed for SENP7 and ORC proteins (Romeo et al., 2015; Maison et al., 2012; Prasanth et al., 2004; Prasanth et al., 2010), we established SUV420H1/H2 double knockout (DKO) HeLa cells, in which H4K20me3 was diminished (Fig. 5A and S5). HP1β signals in H3K9me3-enriched heterochromatin in DKO cells appeared slightly decreased compared to parental cells, although the changes were not always significant in DKO clones (Fig. 5A, 5B and S6A). When HTM-sfGFP was overexpressed in DKO cells, HP1β heterochromatin accumulation was increased but in lesser extent than the parental HeLa cells (Fig. 5C, D and S6B). These results, in one hand, suggest that the endogenous SUV420H2 contributes subtly to HP1 locking probably because its endogenous expression level is not very high. On the other hand, as the increase of HP1β accumulation in H3K9me3-enriched heterochromatin in DKO cells was limited compared to parental cells (Fig. 5D), other functions of SUV420H2 than the HTM-mediated locking, such as adding H4K20me3 may enhance HP1 heterochromatin accumulation.

**Fig. 5.**
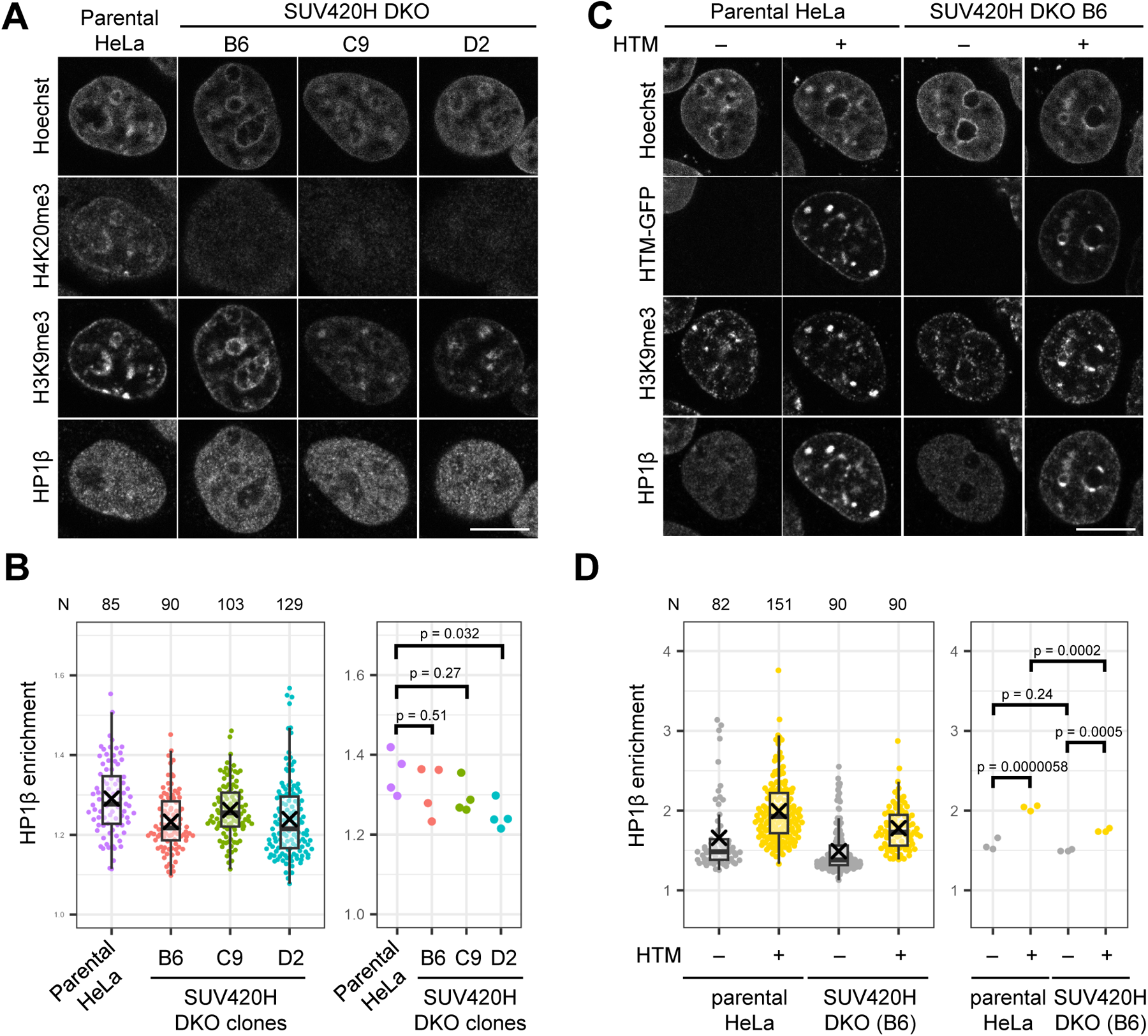
Endogenous SUV420H2 subtly contribute to HP1β locking. (A and B) Slight decrease of HP1β enrichments in H3K9me3-enriched heterochromatin in SUV420H1/H2 double knockout (DKO) HeLa cells. Parental and three DKO HeLa cell lines were stained with antibodies specific to H4K20me3, H3K9me3, and HP1β. (A) Single confocal images of typical nuclei are shown. See Fig. S5A for larger fields of view containing multiple nuclei. (B) Box plots. Average HP1β intensities in the top 10% of H3K9me3 highest intensity pixels were measured and normalized using the mean intensities in entire nuclei. HP1β enrichment in H3K9me3-enriched heterochromatin in individual nuclei from a single experiment is represented as dot and box plots on the left, with the numbers of nuclei (*N*) analyzed indicated above. Mean values of individual experiments in biological quadruplicates are shown on the right. p-values obtained from Dunnett’s test are indicated. Between 83-136 nuclei were analyzed for each cell line in single experiment. (C and D) The effect of HTM-sfGFP on HP1β accumulation in H3K9me3-enriched heterochromatin in DKO cells. Parental HeLa and DKO (clone B6) cells transfected with HTM-sfGFP were stained with antibodies specific to H3K9me3 and HP1β. (C) Single confocal images of typical nuclei. See Fig. S5B for larger fields of view containing multiple nuclei. (D) Box plots ad described in (B). Experiments were performed in triplicate with 52-156 cells analyzed in each line in single experiments. p-values obtained from Tukey-Kramer are indicated. Scale bars: 10 μm.

### HTM V374D-sfGFP can be used for visualizing endogenous HP1 on H3K9me3-enriched heterochromatin

The HTM V374D fragment, which harbors two PxVxL-like motifs for HP1 binding, localized to constitutive heterochromatin in living cells without exhibiting HP1 locking and unlocking effects. Therefore, we thought that HTM V374D could be utilized as a visualization probe for tracking endogenous HP1 on heterochromatin. While GFP-tagged HP1 proteins have been used for labeling heterochromatin and live-cell tracking, their localization to heterochromatin in human cells is less distinct compared to their pronounced highlighting of chromocenters in mouse cells. We compared the localization of HTM V374D-sfGFP, HP1α-sfGFP (C-terminus GFP version), and sfGFP-HP1α (N-terminus GFP version) in mouse A9 and human HeLa cells, which also expressed HaloTag-tagged histone H2B (H2B-Halo). In A9 cells, all three sfGFP-tagged proteins accumulated at chromocenters, identified by H2B-Halo concentration. However, HTM V374D-sfGFP highlighted chromocenters more clearly than HP1α-sfGFP and sfGFP-HP1α (Fig. 6A and S7). In HeLa cells, the differences among the three proteins were more pronounced (Fig. 6B and S7). HTM V374D-sfGFP was enriched in H2B-Halo-enriched regions, whereas HP1α-sfGFP and sfGFP-HP1α were more homogenously distributed throughout the nucleus, often with tiny foci devoid of H2B-Halo, which likely represent PML bodies (Fig. 6B, arrows) (Lehming et al., 1998; Seeler et al., 1998; Hayakawa et al., 2003; Everett et al., 1999). Since HP1 interacts with SP100, a constitutive component of PML bodies, through its CSD (Hayakawa et al., 2003), GFP-tagged HP1 can also target PML bodies via such interactions. As HTM V374D binds to the HP1 CSD, HP1 molecules that are localized to specific compartments via CSD could not be recognized by HTM V374D. Indeed, during mitosis, HTM V374D-sfGFP dissociated from centromeres, to which HP1 binds via CSD, as well as from condensed chromosomes due to histone H3 phosphorylation (Fischle et al., 2005; Hirota et al., 2015), distributing throughout the cytoplasm (Fig. 6). Thus, HTM V374D-sfGFP can be useful to specifically visualize H3K9me3-bound HP1 on interphase heterochromatin.

**Fig. 6.**
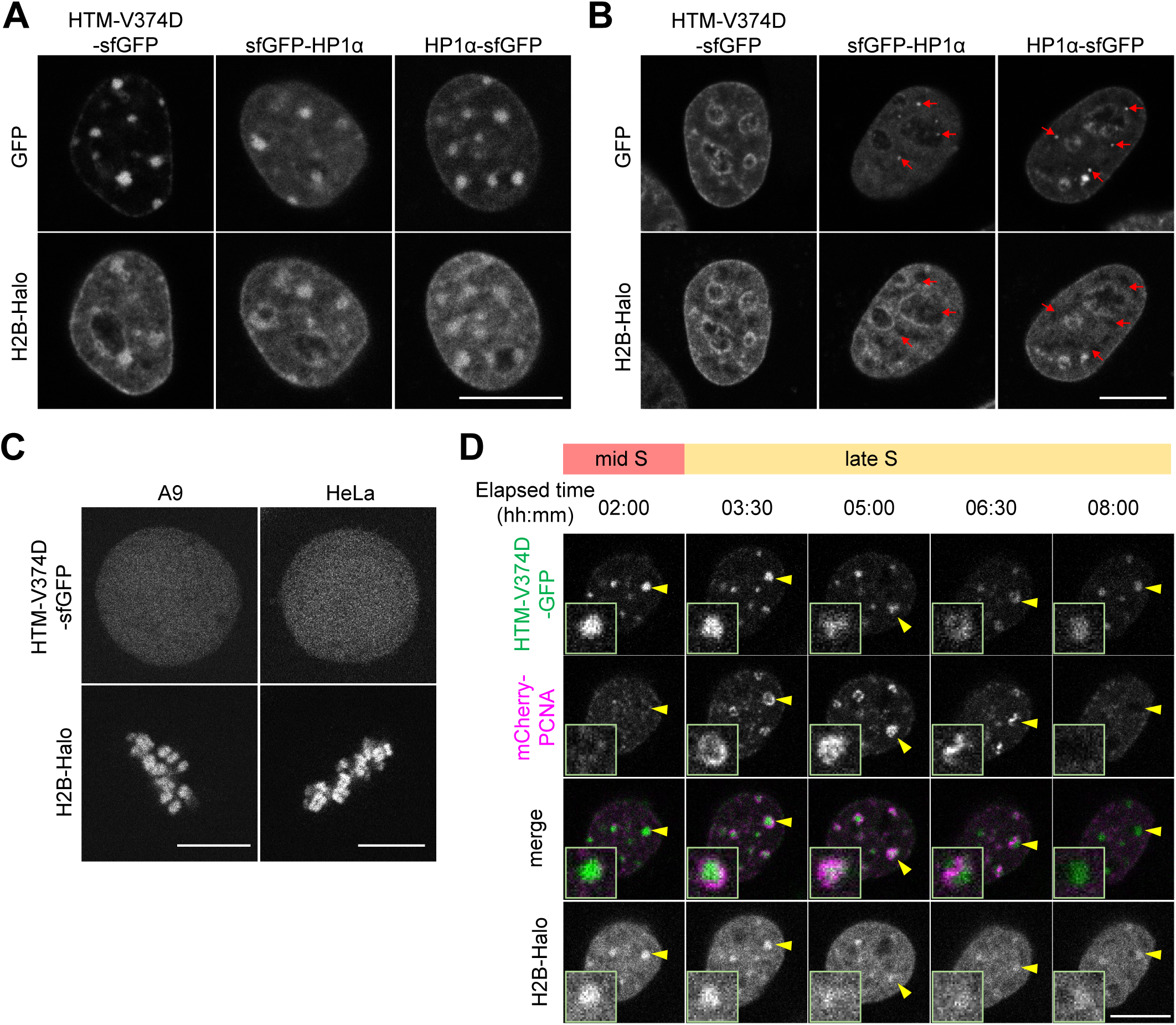
Visualizing HP1-bound heterochromatin in living cells using HTM V374D-sfGFP. (A-B) Distribution of HTM V374D-sfGFP, sfGFP-HP1α, HP1α-sfGFP in interphase nuclei of A9 (A) and HeLa cells (B). H2B-Halo was co-expressed to visualize global distribution of chromatin. Arrows indicate HP1 accumulation in small foci devoid of H2B-Halo. (C) Distribution of HTM V374D-sfGFP throughout the cytoplasm in mitotic cells. (D) Dynamics of HP1-enriched heterochromatin during DNA replication in the late S phase. Yellow arrowheads indicate the same foci, with high-power views shown in insets. Scale bars: 10 μm.

To demonstrate the use of HTM V374D-sfGFP as a live-cell heterochromatin marker, we tracked the dynamics of HTM V374D-sfGFP in chromocenters during the S-phase progression in A9 cells, together with H2B-Halo and mCherry-PCNA to visualize chromatin and DNA replication foci (Chagin et al., 2016; Leonhardt et al., 2000), respectively (Fig. 6D and Movies 1 and 2). Previous studies using fixed cells have reported that DNA replication in heterochromatin begins at the periphery of HP1-rich domains, with replicated DNA moves toward the domain centers (Quivy et al., 2004). However, there are no reports observing the dynamics of HP1-rich domains during replication through live imaging. We observed mCherry-PCNA accumulating around domains enriched in HTM V374D-sfGFP – and so chromatin-bound HP1 – during the late S phase. A detailed view revealed that mCherry-PCNA first accumulated at the periphery of a chromocenter and then expanded towards the center. During this process, the HTM-V374D-sfGFP focus became fuzzy and transformed into a donut-like shape (Fig. 6C, 03:30-05:00; Movie 2). As the mCherry-PCNA signal decreased upon the completion of DNA replication, the HTM-V374D-sfGFP focus reformed into an oval shape (Fig. 6C, 05:00-8:00); Movie 2. This observation suggests that HP1 may temporarily dissociate from heterochromatin during DNA replication, and/or heterochromatin may become decondensed during this process (Chagin et al., 2019; Leonhardt et al., 2000).

## Discussion

### HP1 binding of SUV420H2 HTM

In this study, we identified three HP1 binding motifs within the HTM that are required for efficient heterochromatin localization of SUV420H2. The first and second motifs can function cooperatively, assisted by the zinc-finger-like motif harboring one His and two Cys residues. The full HTM containing the all three motifs, as well as the 29-aa HTM-N peptide containing two PxVxL-like motifs, possesses HP1 locking activity. This stabilizes HP1 on heterochromatin through bivalent binding, as observed in SENP7, Orc3 and PRR14, all of which have multiple HP1 binding motifs (Romeo K 2015, Maison C 2012, Prasanth SG 2004, Prasanth SG 2010, Kiseleva AA 2023). The 22-aa HTM-C peptide, which harbors the third motif with an overlapped PxVxL and PxxVxL stretch, also targets heterochromatin and slightly destabilizes the HP1-chromatin interaction. This is likely due to unlocking, as demonstrated by SENP7’s single HP1 binding motifs accelerating HP1 dissociation (Romeo K 2015). Hence, the stability of HP1 chromatin binding is likely controlled by the balance between different effector proteins harboring single and multiple cooperative HP1 binding modules. The full HTM, containing all three motifs, exhibits the highest HP1 locking effect. However, mutation in the second HP1 binding motif (V374D) abolishes this effect, while the other two motifs remain. Therefore, HP1 locking efficiency depends on the configuration of multiple HP1 binding sites.

Given the strong HP1 locking function in SUV420H2 HTM, a decrease of HP1 locking in SUV420H1/H2 DKO cells could be expected. However, HP1 levels at heterochromatin in DKO cells exhibited only a subtle decrease compared to those in wild-type HeLa cells. The expression level of SUV420H2 in HeLa cells may be too low to induce substantial HP1 locking, and the multiple HP1 binding sites may simply increase its heterochromatin targeting efficiency for H4K20me3 deposition in an HP1-dependent manner. In contrast, H4K20me3 deposited by SUV420H2 appears to enhance the capacity of the HP1 locking, likely through recruiting proteins such as LRWD1/ORCA, which binds to ORC, and DNA methyltransferase 1 (Vermeulen et al., 2010; Ren et al., 2011). A subtle HP1 locking induced by SUV420H2 and/or H4K20me3 may also induce nano-compaction, detectable through energy transfer between two histones (Dupont et al., 2023). It has been reported that HP1β is functionally associated with SUV420H2 and H4K20me3 (Bosch-Presegué et al., 2017), which may be consistent with our finding that the HP1 locking effect induced by HTM-N expression was more pronounced in HP1β compared to HP1α and HP1γ. The detailed molecular mechanism determining how HTM binds to different HP1 subtypes through amino acids surrounding the PxVxL motifs (Canzio et al., 2014) remains to be elucidated.

### Visualizing HP1-bound heterochromatin using HTM V374D

As HTM V374D does not induce HP1 locking nor unlocking, this fragment can be applicable for visualizing heterochromatin marked by HP1. Although HP1 could be directly visualized by tagging with the fluorescent proteins, GFP-tagged HP1 does not always exhibit heterochromatin localization in human cells, whereas it is concentrated in chromocenters in mouse cells (Fig. 6 and S7) (Dialynas et al., 2007), suggesting that the bulky protein tag may interfere with the binding to H3K9me3 and/or effector proteins. The binding of HTM V374D probe to HP1 CSD could also block the binding of effector proteins. However, under low expression levels, potential blocking effects may be minimal because the dynamics of HP1 remains unchanged when HTM V374D is expressed. Indeed, we have demonstrated that HTM V374D can be used for tracking DNA replication at chromocenters during the mid to late S phase, in combination with mCherry-PCNA. We observed that HTM V374D probe became less concentrated, either by dissociation or massive decondensation (Chagin et al., 2019; Leonhardt et al., 2000), when replication occurs from the surface of chromocenters. To monitor H3K9me3 in living cells, several probes based on HP1 CD have been developed (Sánchez et al, 2019; Villaseñor et al., 2020; Sasaki et al., 2022). The binding of these HP1 CD-based probes to H3K9me3 can be influenced by H3S10 phosphorylation, similar to HTM V374D, which dissociates from mitotic chromosomes. The HTM V374D probe, as reported here, offers an additional option for tracking HP1-bound H3K9me3-rich heterochromatin in living cells.

## Materials and methods

### Cells and transfection

HeLa (obtained from Peter R. Cook at Oxford University; Pombo et al., 1999), A9 (obtained from Nobuo Takagi at Hokkaido University), and NIH3T3 cells (obtained from Nobuo Takagi at Hokkaido University) were grown in Dulbecco’s modified Eagle’s medium (DMEM), high-glucose (Nacalai Tesque) containing 10% fetal bovine serum (FBS) (Gibco, Thermo Fisher Scientific) and 1% L-glutamine–penicillin–streptomycin solution (GPS; Sigma-Aldrich) at 37°C in a 5% CO_2_ atmosphere. For transfection, FuGENE HD (Promega) was used according to the manufacturer’s instructions. Briefly, 2 µg DNA was mixed with 6 µL of FuGENE HD in 100 µL of Opti-MEM (Thermo Fisher Scientific) and incubated at RT for 10 min before being added to HeLa cells grown in 35-mm glass-bottomed dishes (AGC Technology Solutions) at 40–70% confluency. To obtain stably expressing cells, 1.8 µg PB533-or PB510-based PiggyBac plasmid (System Biosciences) and 0.2 µg transposase expression vector (System Biosciences) were used, and cells were selected in 1 mg/mL G418 (Nacalai Tesque). Cells stably expressing sfGFP fusion proteins at similar levels among different mutants were collected using a cell sorter (SH800; Sony).

### Plasmid construction

The HaloTag-tagged KDM4D expression vector (FHC06842; Promega), the PB533-based H2B-Halo expression vector (Uchino et al., 2021), and the PB533-based mCherry-PCNA expression vector (Uchino et al., 2021) were reported previously. For transient expression in mammalian cells, psfGFP-N1 (Addgene 54737) and psfCherry-C1 (Kono et al., 2022) were used. To construct SUV420H2 deletion mutants, the HaloTag-tagged SUV420H2 expression vector (FHC26822; Promega) was used as a template for PCR amplification of the target regions. The primer used for PCR in this study are listed in Supplementary Table S2. The amplified DNA fragments were then cloned into psfGFP-N1 linearized by EcoRI and BamHI digestion, using the In-Fusion ^®^ HD Cloning Kit (Z9648N; TaKaRa).

To construct psfCherry-HP1α, HP1α coding region was amplified by PCR using the GFP-HP1α expression vector provided by C. Obuse (Nozawa et al., 2010) as a template. The amplified DNA fragment was cloned into psfCherry-C1 linearized by EcoRI digestion. To introduce point mutations, inverse PCR was conducted (Uchino et al., 2021). Some coding sequences with amino acid substitutions were amplified using a plasmid that already harbored another mutation. For establishing stably expressing cells, Piggy Bac system was used. The fragment coding HTM mutants-sfGFP in psfGFP-N1 was exercised by EcoRI and NotI digestion and ligated into the PB533 vector digested with EcoRI and NotI. The psfGFP-352-(GGGGS)×3-380 was constructed by insertion of a flexible linker using primers (pSUV420H2[352-]_sfGFP_InF_s, pSUV420H2[-380]_sfGFP_InF_as, LINK.AMP5T, LINK.AMP3T, 370-GSLINK_s, 369-GSLINK_as) as described previously (Sato et al., 2013). The nucleotide sequence of all the inserts was verified.

### Live cell microscopy

For Fig. 1, 3A, 3B, S1C, and S1D, A9 cells (Fig. 1, 3A, 3B, and S1D) or those stably expressing HTM-sfGFP (Fig. S1C) were plated in a 24 well glass-bottom plate (AGC Technology Solutions) at a cell density of 0.6-1.2 × 10^5^ cells/well. The following day, cells were transfected with plasmids to express sfGFP-tagged SUV420H2 mutants (Fig. 1, 3A, 3B, and S1D) or Halo-KDM4D (Fig. S1C). One day post-transfection, the medium was replaced with FluoroBrite (Thermo Fisher Scientific) containing 10% FBS and 1% GPS. To detect Halo-KDM4D, cells were incubated with HaloTag TMR Ligand (Promega) at a final concentration of 100 nM for 30 min incubation, before replacing the medium. The cell culture plates were placed on a heated stage (Tokai Hit) at 37°C under 5% CO_2_ regulated by a CO_2_ control system (Tokken) on a confocal microscope (FV-1000; Olympus) operated by built-in software (Fluoview ver. 4.2) with a PlanSApo 60× (NA 1.40) oil-immersion objective lens. Images were acquired using the line-sequential imaging mode (2.0% 488-nm laser transmission; 1024 × 1024 pixels; pinhole 200 μm; 2× zoom for HeLa cells or 3× zoom for A9 cells; 2-line Kalman filtration) with a 405/488/543/633 dichromic mirror and a 505-525 emission filter. Image analysis was performed using the NIS-elements Analysis software ver. 5.1 (Nikon). After background subtraction, the chromocenters and the entire nucleus were both selected by Magic Wand tool to measure the fluorescence intensities in the selected areas. Heterochromatin enrichment ratios were calculated by dividing the mean intensity of chromocenters by that of the nucleus.

For Fig. 6, HeLa and A9 cells were plated in a 24-well glass bottom plate (AGC Technology Solutions) at a cell density of 0.6-1.2 × 10^5^ cells/well. The next day, cells were transfected with plasmids to co-express HTM V374D-sfGFP and H2B-Halo. Two days post-transfection, HaloTag TMR Ligand (Promega) was added at a final concentration of 100 nM, followed by a 30-min incubation, after which the medium was replaced with FluoroBrite (Thermo Fisher Scientific) containing 10% FBS and 1% GPS. For Fig. 6C, HeLa and A9 cells stably expressing HTM V374D-sfGFP, mCherry-PCNA, and H2B-Halo, plated in a 24-well glass-bottom plate were incubated with Janelia Fluor 646 HaloTag Ligand (Promega) for 30 min, and the medium was replaced with FluoroBrite containing 10% FBS and 1% GPS. The cell culture plates were set on a heated stage at 37°C under 5% CO_2_ (Tokai Hit) on a confocal microscope (A1R; Nikon) operated by NIS Elements ver. 5.21.00 (Nikon) with an Apo TIRF 60× (NA 1.49) oil-immersion objective lens. Images were collected using the line-sequential imaging mode (1024 × 1024 pixels; pinhole 24.27 μm; 2× zoom for HeLa or 3× zoom for A9; 2-line Kalman filtration) with two or three laser lines (0.4-1.0% 488-nm laser transmission; 1.0% 561-nm laser transmission; 0.2% 640-nm laser transmission) with a 405/488/543/633 dichromic mirror and 525/50, 595/50, and 700/75 emission filters.

### Immunofluorescence

Antibodies and staining conditions used in this study are listed in Supplementary Table S2. For Fig. S1B, HeLa, NIH3T3, and A9 cells grown on a 24-well glass-bottom plate were transfected with expression plasmids of HTM-sfGFP and its derivatives. One day post-transfection, cells were fixed with 4% paraformaldehyde (PFA; Electron Microscopy Sciences) in 250 mM HEPES-NaOH (pH 7.4) for 5 min at room temperature, washed with Dulbecco’s Phosphate Buffered Saline, calcium- and magnesium-free (PBS; Fujifilm Wako Chemicals), and permeabilized using 1% Triton X-100 (Nacalai Tesque) for 20 min with gentle shaking. Cells were then incubated in Blocking-One P solution (Nacalai Tesque) for 20 min with gentle shaking and then stained with Hoechst 33342 (1 µg/mL) and mouse monoclonal antibody directed against H3K9me3 directly-labeled with Cy3 (2 μg/mL; Chandra et al., 2012) in 10% Blocking-One P in PBS for 3 h at room temperature, before washing with PBS three times. Fluorescence images were collected using a spinning disk confocal microscope system (CSU-W1; Yokogawa and Ti-E; Nikon) with a PlanApo VC 100× (NA 1.40) oil-immersion objective lens, a 405/488/561/640 dichroic mirror, and 450/50, 525/50, 595/50, and 700/75 emission filters, equipped with an electron-multiplying charge-coupled device (iXon+; Andor) and 405-, 488-, 561-, and 640-nm laser lines (LU-N4; Nikon).

For Fig. S5A, parental HeLa and SUV420H1/2 DKO cells were plated on a 24-well glass-bottom plate. The following day, cells were fixed with 4% PFA in 250 mM HEPES-NaOH (pH 7.4) with 0.1% TritonX-100 for 5 min at room temperature, then permeabilized and blocked as described above, before staining with Hoechst 33342 (1 µg/mL) and fluorescent dye-labeled mouse monoclonal antibodies specific to H4K20me3 (Cy3), H4K20me2 (Alexa488) and H4K20me1 (Cy5) (4 μg/mL for each antibody) in 10% Blocking-One P in PBS at 4°C overnight (Hayashi-Takanaka et al., 2015). Cells were then washed with PBS three times. The samples were set on a confocal microscope (A1R; Nikon) operated by NIS Elements ver. 5.21.00 (Nikon) with a PlanApo λ 100× (NA 1.45) oil-immersion objective lens, a 405/488/561/640 dichroic mirror, and 450/50, 525/50, 595/50, and 700/75 emission filters. Images were collected using line-sequential imaging mode (1024 × 1024 pixels; pinhole 255.43 μm; 2× zoom; 2-line Kalman filtration) with four laser lines (LU-N4; Nikon; 0.1% 405-nm laser transmission; 0.1% 488-nm laser transmission; 0.1% 561-nm laser transmission; and 0.1% 640-nm laser transmission). For quantitative analysis, individual nuclei were defined using magic wand tool in NIS Elements ver. 5.30.02.

### HP1 locking assay by immunofluorescence

Cell were plated in a 35-mm glass-bottom dish a day before fixation (total 4-5 × 10^5^ cells; Fig. 4A, 5A, 5B, S3A, S4A, and S6A; cells expressing sfGFP-tagged protein and the parental cells were mixed at a 1:1 ratio) or a day before transfection (2–2.5 × 10^5^ cells; Fig. 5C and S6B). One day after cell plating or transfection, the cells were fixed, permeabilized, blocked, and stained with the primary and the secondary antibodies (Supplementary Table S2), as described above. Fluorescence images were acquired using a confocal microscope (A1R; Nikon) operated by NIS Elements ver. 5.21.00 (Nikon) with an Apo TIRF 60× (NA 1.49) oil-immersion objective lens, a 405/488/561/640 dichroic mirror, and 450/50, 525/50, 595/50, 700/75 emission filters, using the line-sequential imaging mode (1024 × 1024 pixels; pinhole 26.82 μm; 2× zoom; 2-line Kalman filtration) with four laser lines (0.3-0.6% 405-nm laser transmission; 0.3-0.4% 488-nm laser transmission; 0.1%-4% 561-nm laser transmission; 0.3-0.8% 640-nm laser transmission).

For quantitative analysis for Fig. 4B, 5B and 5D, individual nuclei were defined using the Magic Wand tool in NIS Elements ver. 5.30.02 and minimum rectangle areas containing single nuclei were cropped for exporting as TIF files. The intensity in each pixel was measured using “TIFF” package in R (https://www.r-project.org/). To compare the changes in the enrichment ratio of HP1β in the H3K9me3-enriched domain, the pixels showing the top 10% highest intensity in the H3K9me3 channel were selected and then the mean intensity of HP1β in these H3K9me3-top10% pixels was divided by that in the whole nucleus for normalization. To compare the changes of HP1α, HP1β and HP1γ signals by HTM-N-sfGFP expression, the mean intensity of HP1 signals was divided by the mean value of the mean intensities of HP1 in control cells.

### HP1 locking assay by fluorescence recovery after photobleaching (FRAP)

A9 cells plated in a 24-well glass bottom plate were co-transfected with transient expression plasmids of HTM-sfGFP, HTM-N-sfGFP, HTM-C-sfGFP, or HTM V374D-sfGFP, along with the sfCherry-HP1α expression plasmid. The next day, the medium was replaced with FluoroBrite (Thermo Fisher Scientific) containing 10% FBS and 1% GPS, before a dish was placed on a heated stage (Tokai Hit) at 37°C under 5% CO_2_ regulated by a CO_2_ control system (Tokken) on a confocal microscope (FV-1000; Olympus), operated by built-in software (Fluoview ver. 4.2) with a PlanSApo 60× (NA 1.40) oil-immersion objective lens. For the FRAP experiment, twenty images were collected (10% 543-nm laser transmission; 128×32 pixels; pinhole 800 μm; 12× zoom). A 1.38 μm diameter circular area was bleached (100% transmission for 458-, 488-, 515-, and 543-nm laser lines; duration 364.32 ms), followed by the consecutive collection of another 80 images. Fluorescence intensity measurements were performed using NIS Elements ver. 5.30.02. The net intensities of the bleached and unbleached areas were calculated by subtracting the background intensity outside nuclei in each time frame. To calculate relative intensities to the initial intensity of the bleached area, the global loss of fluorescence was normalized by dividing the intensity in bleached area by that in unbleached area, before normalizing to the average intensity of the pre-bleach images.

### Double knockout of SUV420H1/H2

Lacking access to a highly specific antibody for SUV420H2 knockout validation, we decided to establish SUV420H1/H2 double knockout (DKO) cells. These cells were reported to have almost no H4K20me3 and H4K20me2, and have greater level of H4K20me1 (Schotta et al., 2008). To establish DKO cells of SUV420H1 and SUV420H2, the CRISPR/Cas9 system (pX330, Addgene 42230; pKN7, Addgene DU70250; and pX459, Addgene 62988) was used. Plasmid construction followed the “Target Sequence Cloning Protocol” provided by Feng Zhang lab (https://www.addgene.org/crispr/zhang/). The gRNAs were designed using the “CRISPR Finder” (Wellcome Sanger Institute Genome Editing; https://wge.stemcell.sanger.ac.uk/find_crisprs), targeting the first exons and SET domains, considering several known splicing isoforms. The sequences of the primers used for plasmid construction are summarized in supplemental Table S1. HeLa cells were plated in a 6-well plate at a density of 2.4 × 10^5^ cells/well. The following day, cells were transfected with plasmids for KO (pX330 and pKN7 for the first exons; and pX459 for SET domains) using Lipofectamin 2000 according to the manufacturer’s instruction. One and three days post-transfection, the medium was replaced to DMEM containing 1 μg/mL puromycin. For single cell cloning, puromycin-resistant cells were seeded at a density of 50-100 cells in a 10 cm dish. After a week, single colonies were picked and transferred to wells in two 96-well plates; one with a plastic bottom for stock and another with a glass bottom for immunofluorescence. To screen and validate DKO cells, immunofluorescence was performed using the fluorescent dye directly-labeled mouse monoclonal antibodies against H4K20me3 (Cy3), H4K20me2 (Alexa488), and H4K20me1 (Cy5). DKO cells were generated by three steps. First, the first exons of SUV420H1 and SUV420H2 were simultaneously targeted. As H4K20me2 and H4K20me3 signals were still observed, the SET domain of SUV420H1 was targeted next. In one clone, post-immunofluorescence screening, H4K20me2 signals were diminished to background level but some H4K20me3 signals remained. The SET domain of SUV420H2 was then targeted. Three clones (B6, C9, and D2) showed no signals for both H4K20me2 and H4K20me3 (Fig. S5).

### Time-lapse observation of the HP1 dynamics during the late S phase

A9 cells stably expressing sfGFP-HTM-C, mCherry-PCNA, and H2B-Halo, were plated on a 35-mm glass-bottom dish (AGC Technology Solutions). The next day, Janelia Fluor 646^®^ HaloTag Ligand^®^ was added to the medium at a final concentration of 100 nM. After a 30-min incubation, the medium was replaced with FluoroBrite (Thermo Fisher Scientific) containing 10% FBS and 1% GPS. The dish was placed on a spinning disk confocal microscope system (CSU-W1; Yokogawa and Ti-E; Nikon) equipped with a PlanApo VC 100× (NA 1.4) oil-immersion objective lens (with Type F immersion oil; MXA22168; Nikon) and integrated with a culture system (Tokai Hit) maintained at 37°C under 5% CO_2_. Fluorescence images were captured using NIS Elements ver.5.11.03 (Nikon), operated with an LDI-7 Laser Diode Illuminator (Chroma Technologies Japan; 10% 470-nm laser transmission; 20% 555-nm laser transmission; 50% 640-nm laser transmission) was used with a 405/470/555/640 dichroic mirror, a 520/60, a 600/50, and a 690/50 emission filters, and an EM charge-coupled device (iXon+, Andor; gain 300; exposure time 470 nm: 1s, 555 nm: 1s, 640 nm: 500 ms). Nine different focal planes were imaged at 0.5 µm intervals every 5 min.

### Immunoprecipitation and western blotting

HeLa cells expressing the sfGFP fusion proteins (4.4 × 10^6^ cells) were plated on a 10-cm cell culture dish (Greiner). The next day, cells were collected using a cell lifter (Corning), washed with ice-cold PBS (Takara), and resuspended in 500 µL of ice-cold Lysis buffer [1 M NaCl, HEPES-NaOH (pH 7.4, Nacalai Tesque), 300 mM sucrose (Nacalai Tesque), 0.1% Triton X-100 (Nacalai Tesque), 1 mM MgCl2 (Sigma Aldrich), 1 mM EGTA (Nacalai Tesque)]. Small aliquots (20 µL) were kept as “Whole cell” fractions for immunoblotting. After centrifugation (20,000 ×*g*, 20 min, 4°C), the supernatants (470 µL) were collected and 625 U of Benzonase (Novagen) was added to each sample to digest nucleic acids, thereby eliminating DNA/RNA-mediated interactions. The pellets were resuspended in the original volumes of Lysis buffer for “Insoluble pellet” fractions. After a 30-min incubation on ice, the supernatants (450 µL) were transferred to new tubes and mixed with an equal volume (450 µL) of NaCl-free Dilution Buffer [HEPES-NaOH (pH 7.4), 300 mM Sucrose, 0.1% Triton X-100, 1 mM MgCl2, 1 mM EGTA] to lower the salt concentration to 500 mM for immunoprecipitation. Following centrifugation (20,000 ×*g*, 20 min, 4°C), the supernatants were collected, and 20 µL aliquots were kept as “IP input” fractions for immunoblotting. The remaining supernatants were mixed with GFP-Trap magnetic beads (8 µL of slurry per sample; Chromotek, gtma-20) prewashed with IP Buffer [500 mM NaCl, HEPES-NaOH (pH 7.4), 300 mM Sucrose, 0.1% Triton X-100, 1 mM MgCl2, 1 mM EGTA]. The mixtures were incubated at 4°C overnight with rotation. After collecting the beads using a magnetic stand (Thermo Fisher Scientific), supernatants were transferred to a new tube for “Unbound” samples. The beads were washed with ice-cold IP Buffer and then resupended in 20 µL of IP Buffer (×44 concentrated compared to the input). To adjust the concentration of “Whole cell” and “Insoluble pellet” fractions to that of “IP input”, an equal volume of NaCl-free Dilution Buffer was added. Samples for immunoblotting were mixed with 2× Sample-Loading Buffer [125 mM Tris-HCl, pH 6.8, 20% glycerol (Fujifilm Wako Chemicals), 4% sodium dodecyl sulfate (SDS; Fujifilm Wako Chemicals), 0.01% bromophenol blue (Fujifilm Wako Chemicals), and 10% dithiothreitol (Fujifilm Wako Chemicals)] and heated at 95°C for 10 min. Then, 5 μL of each sample was separated on 15% polyacrylamide gels (SuperSep™ Ace, 17 well pre-cast; Fujifilm Wako Chemicals) and transferred to FluoroTrans W PVDF Transfer Membranes (Pall; 90 minutes; 170 mA constant for a 9 cm × 9 cm membrane) using EzFastBlot (Atto) as a transfer buffer. The membranes were blocked with Blocking One (Nacalai Tesque) for 30 min with gentle shaking. After washing with TBST [20 mM Tris-HCl, pH 8.0, 150 mM NaCl, 0.02% Tween 20], the membranes were incubated with the primary antibody, i.e., rabbit monoclonal anti-HP1α (1:10,000; Abcam; ab109028), rabbit monoclonal anti-HP1β (1:1,000; Cell Signaling Technology; D2F2, #8676), rabbit polyclonal anti-HP1γ (1:1,000; Cell Signaling Technology; #2619), and rabbit polyclonal anti-GFP (1:2,000; MBL; No.598), in Can-GetSignal® Solution 1 (Toyobo) for 2 h at room temperature. After washing the membranes with TBST three times, they were incubated with horseradish peroxide-conjugated goat anti-mouse or anti-rabbit IgG (H+L) (1:10,000; Jackson ImmunoResearch) in Can-Get-Signal® Solution 2 (Toyobo) for 1 h at room temperature. The membranes were then washed with TBST three times. Western Lightning® Plus-ECL reagent (PerkinElmer) was used for chemiluminescence detection using a gel imaging system (LuminoGraph II, Atto).

### Statistical analysis and data visualization

For statistical analysis, Dunnett’s, Tukey-Kramer, and Student’s t-tests were performed using the lawstat (for Student’s t-test) and multcomp (for Dunnett’s and Tukey-Kramer tests) package in R software (version 4.2.3; https://www.r-project.org/). Line graphs and box plots were drawn using R.

### Amino acid alignment

The amino acid alignments in Fig. S2 were aligned using MAFFT version 7 on the web (https://mafft.cbrc.jp/alignment/server/index.html), employing L-ins-I method. The accession numbers used were: *Homo sapiens*, CCDS12922.1; *Mus musculus*, CCDS20743.2; *Phascolarctos cinereus*, XP_020859355.1; Ornithorhynchus anatinus, XP_028930021.1; *Alligator mississippiensis*, XP_014459640.1; *Chrysemys picta*, XP_023969116.1; *Anolis carolinensis*, XP_016851995.1; *Gallus gallus*, XP_040550951.1.

## Acknowledgements

The authors are grateful to Harumi Ueno (Tokyo Tech) for constructing some plasmids, Haruka Oda and Daiki Maejima (Tokyo Tech) for instructing data analysis using R, Koji Nagao (Osaka Univ) and Hidenori Nishihara (Tokyo Tech and Kindai Univ) for predicting potential HP1 binding motifs and instructing multiple alignment of amino acid sequences, Tetsuya Handa (Tokyo Tech) for instructing basic experimental techniques, Cristina M. Cardoso (TU Darmstadt) and Heinrich Leonhardt (LMU Munich), members of Kimura lab for helpful discussion and suggestions, and the Center for Integrative Biosciences and the Biomaterials Analysis Division, Open Facility Center at the Tokyo Institute of Technology for DNA sequencing.

This work was supported by Japan Society for the Promotion of Science KAKENHI (JP17H01417 and JP21H04764 to H. Kimura), Japan Science and Technology Agency CREST (JPMJCR20S6 to Y. Sato and JPMJCR16G1 to H. Kimura), and Japan Agency for Medical Research and Development (AMED) Basis for Supporting Innovative Drug Discovery and Life Science Research (BINDS) (JP23ama121020 to H. Kimura).

## Author contributions

Y.S. and H.K. conceived the study. M.N. and A.A. performed experiments. M.N. analyzed the data. M.N., Y.S., and H.K. wrote the manuscript.

## Supplementary Figure legends

**Supplementary Fig. S1.**
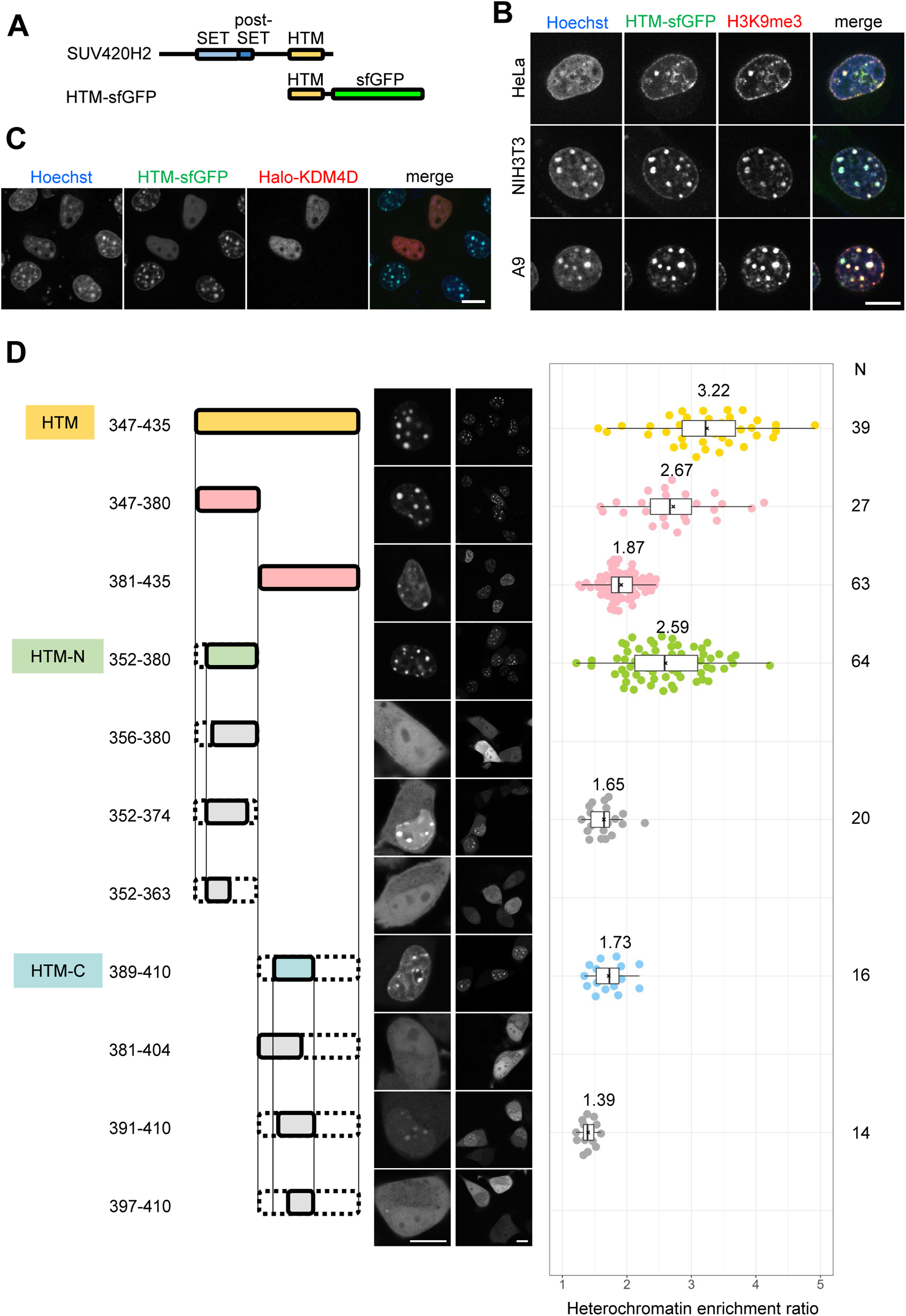
Regions required for heterochromatin localization within SUV420H2 heterochromatic target domain (HTM) dependent on H3K9me3. (A) Schematic representation of SUV420H2 domains. (B) Immunofluorescence analysis of cells expressing HTM-sfGFP. HeLa, NIH3T3, and A9 cells expressing HTM-sfGFP were fixed, stained with an H3K9me3-specific antibody, and counterstained with Hoechst 33342. Single confocal sections are shown. (C) Redistribution of HTM-sfGFP upon H3K9 demethylation induced by Halo-KDM4D expression in A9 cells. Cells stably expressing HTM-sfGFP were transfected with Halo-KDM4D and the next day cells were stained with JF646 HaloTag ligand and Hoechst 33342, and then confocal images were collected. (D) Localization of SUV420H2 HTM sub-fragments. (left) A schematic illustration of the deletion mutants is shown. (middle) Localization patterns of the mutants tagged with sfGFP. Both the high- and low-power views are indicated. (right) Heterochromatin accumulation ratios of the deletion mutants are presented as box plots, indicating the top and bottom 25% with median (thick bar with values) and average (x). The numbers of analyzed nuclei are indicated on the right. Plots for HTM (347-435), HTM-N (352-380), and HTM-C (389-410) are reproductions of those in Fig. 1C. Scale bars: 10 μm.

**Supplementary Fig. S2.**
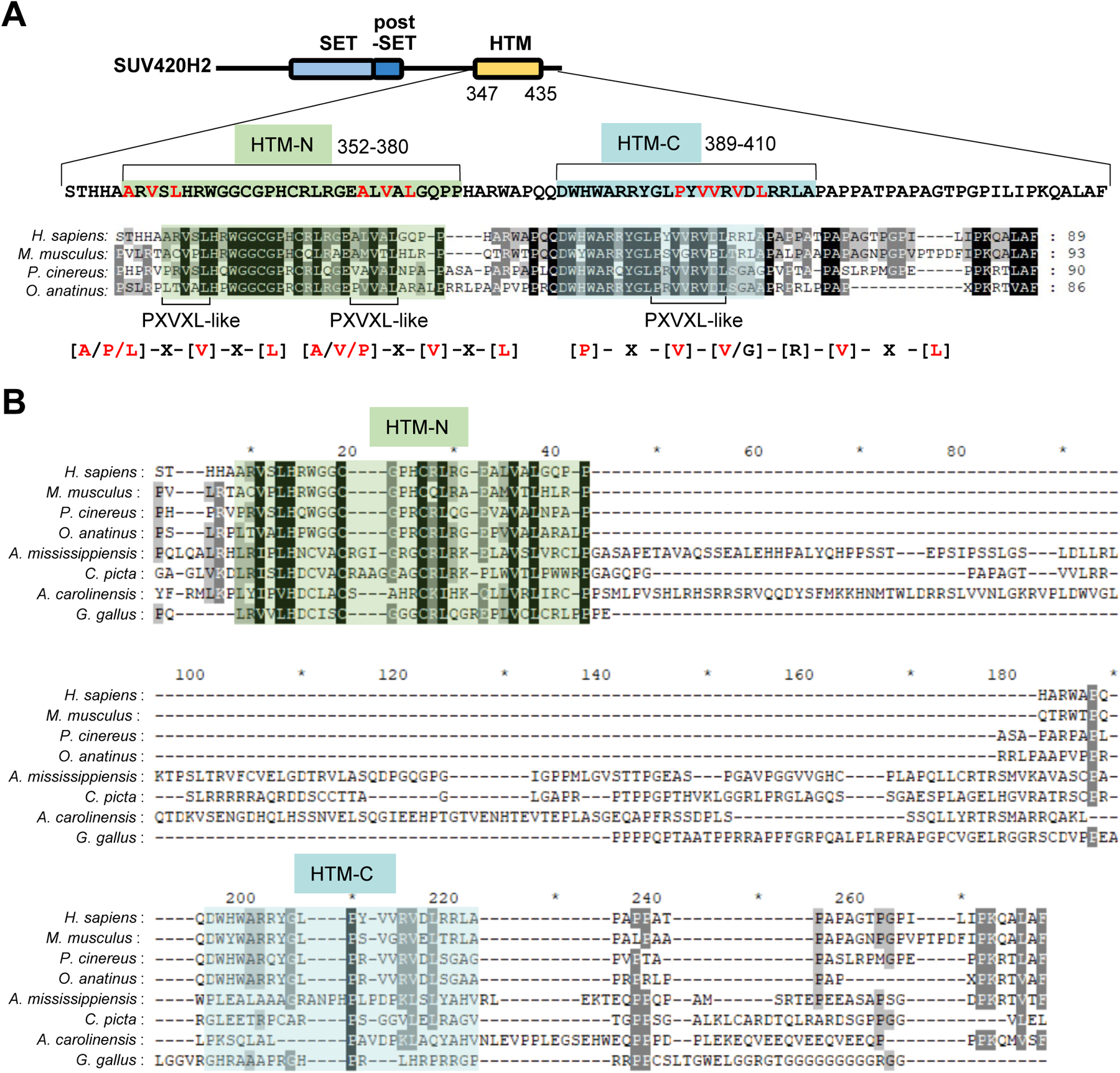
Amino acid sequence alignments. The alignments were generated using MAFFT with L-ins-I protocol. Black, dark gray, and light gray indicate 100%, >80%, and >60% conservation, respectively. Similar groups are considered when counting conserved amino acids. Green- and blue-shaded boxes highlight the regions of HTM-N and HTM-C, respectively. (A) Conserved residues among mammals including *Homo sapiens*, *Mus musculus*, *Phascolarctos cinereus*, and *Ornithorhynchus anatinus*. (B) Conserved residues among amniota, including *Homo sapiens*, *Mus musculus*, *Phascolarctos cinereus*, *Ornithorhynchus anatinus*, *Alligator mississippiensis*, *Chrysemys picta*, *Anolis carolinensis*, and *Gallus gallus*. The HTM-N region is conserved throughout amniota, whereas the HTM-C region is conserved only among mammals.

**Supplementary Fig. S3.**
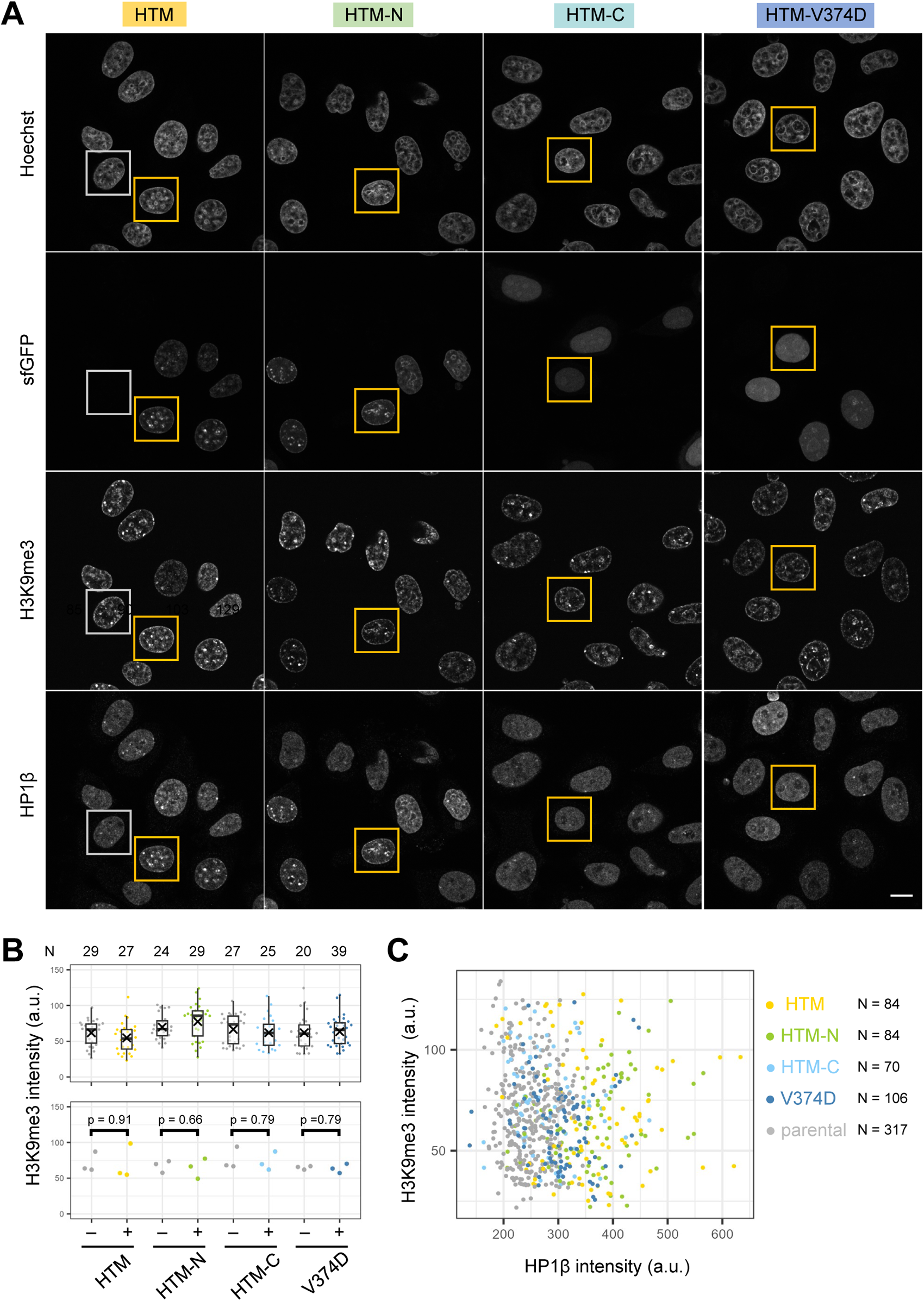
Effects of SUV420H2 HTM expression on HP1β accumulation in H3K9me3-enriched heterochromatin. (A) Views containing multiple nuclei corresponding to Fig. 4A. Enlarged images of nuclei boxed in yellow (sfGFP-tagged protein-expressed) and white (non-expressed) are shown in Fig. 4A. (B and C) H3K9me3 levels were not altered by the expression of HTM and its mutants. (B) H3K9me3 intensities (arbitrary units; a.u.) in individual nuclei from a single experiment with the number of nuclei (*N*) analyzed (top) and mean intensities from single experiments in biological triplicates (bottom) are plotted. p-values obtained from Student’s t-test (paired, two-tailed) are indicated. (C) Scatter plots representing the relationship between H3K9me3 and HP1β intensities, showing no correlation. The number of analyzed nuclei (*N*) is indicated on the right. Scale bar: 10 μm.

**Supplementary Fig. S4.**
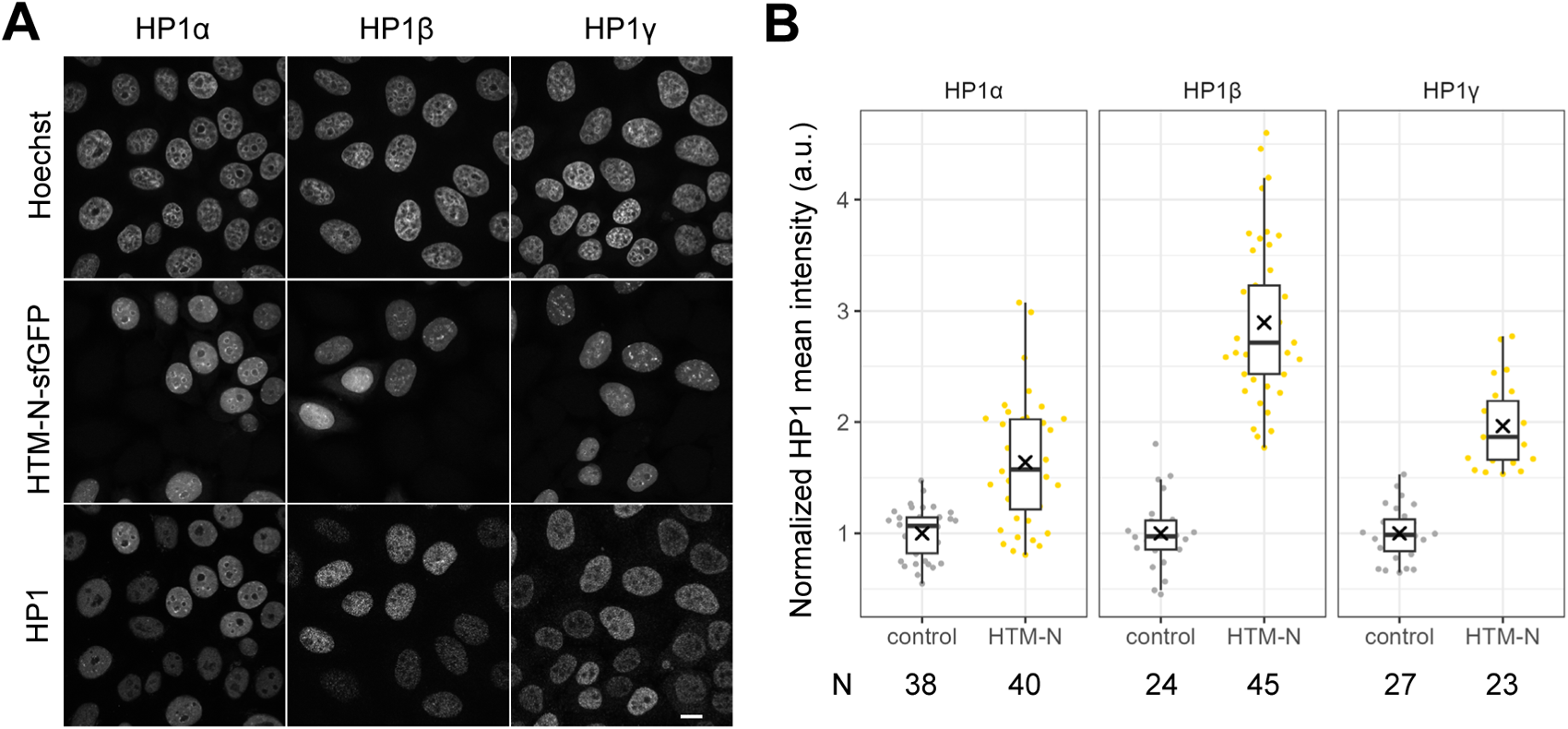
Overexpression of HTM-N-sfGFP results in the increased intensity of all HP1 isoforms. HeLa cells expressing HTM-N-sfGFP were co-plated with parental HeLa cells for immunostaining with antibodies specific to HP1α, HP1β, and HP1γ. DNA was counterstained with Hoechst 33342. (A) Single confocal sections are shown. (B) Boxplots display the mean intensities of HP1 signals with the number of analyzed nuclei (*N*). Scale bar: 10 μm.

**Supplementary Fig. S5.**
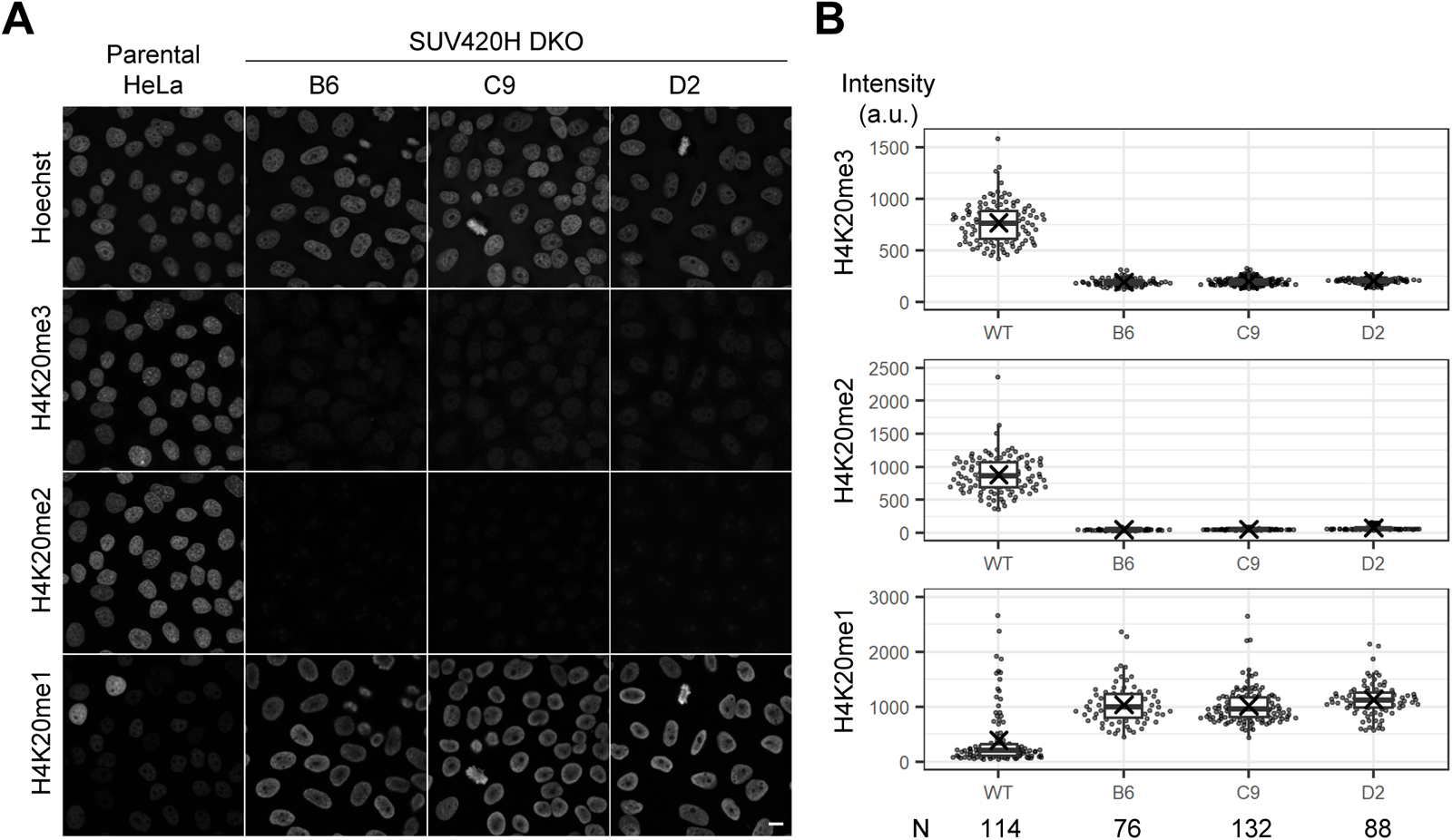
Validation of SUV420H1/H2 double knockout (DKO) cell lines. Levels of H4K20me1, me2, and me3 in parental and SUV420H1/H2 DKO HeLa cells were analyzed by immunofluorescence using specific antibodies conjugated with different fluorescent dyes. (A) Single confocal sections are shown. (B) Boxplots indicate the mean intensities (arbitrary units; a.u.) in individual nuclei. The number of analyzed nuclei is indicated at the bottom. Compared to parental HeLa cells, both H4K20me2 and H4K20me3 levels were diminished to background levels in DKO clones, while H4K20me1 levels were increased. Scale bar: 10 μm.

**Supplementary Fig. S6.**
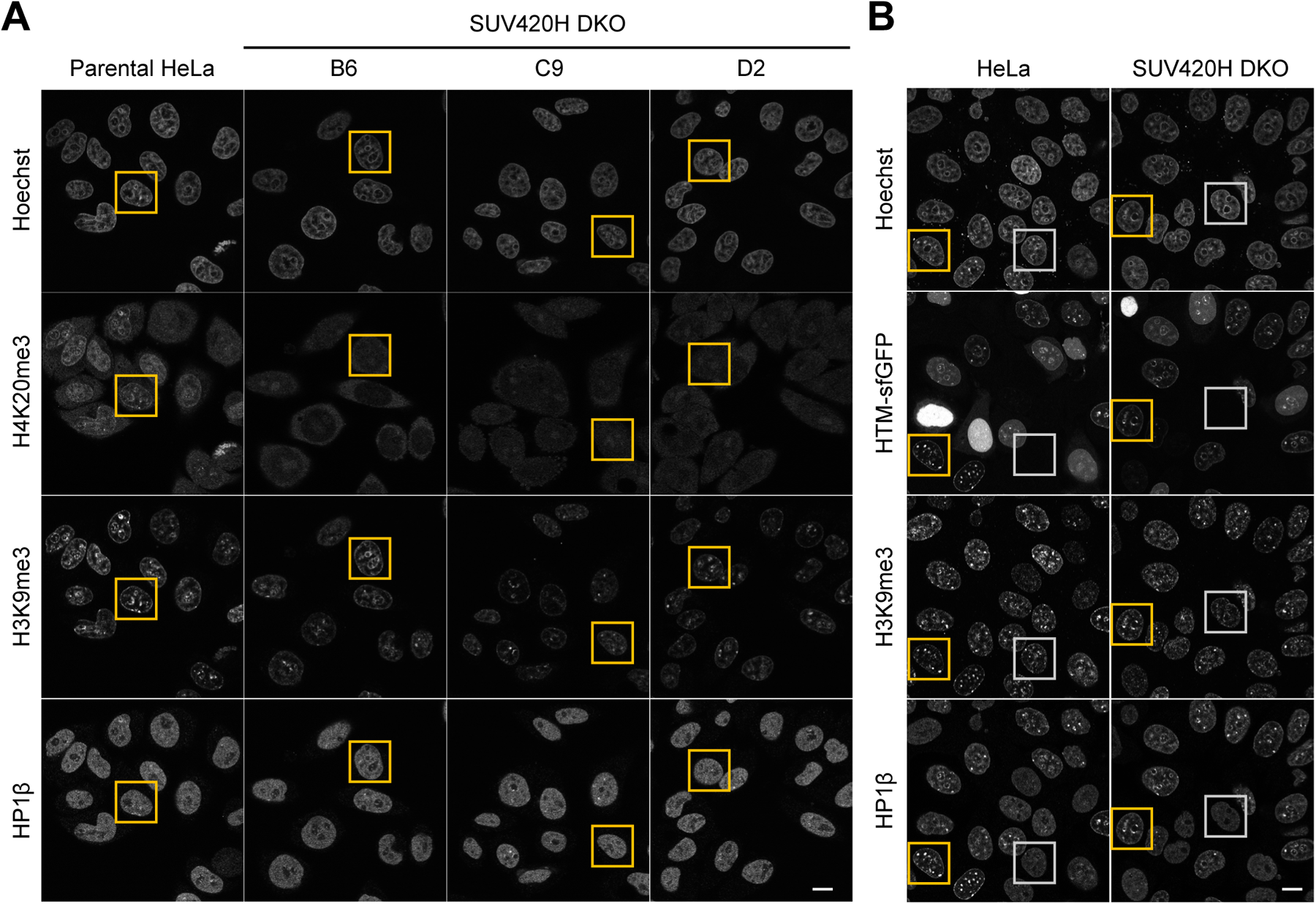
Effects of SUV420H1/H2 DKO and HTM expression in DKO cells on HP1β accumulation in H3K9me3-enriched heterochromatin. (A) Views containing multiple nuclei corresponding to Fig. 5A. Enlarged images of nuclei boxed in yellow are shown in Fig. 5A. (B) Views containing multiple nuclei corresponding to Fig. 5C. Enlarged images of nuclei boxed in yellow (sfGFP-tagged protein-expressed) and white (non-expressed) are shown in Fig. 5C. Scale bars: 10 μm.

**Supplementary Fig. S7.**
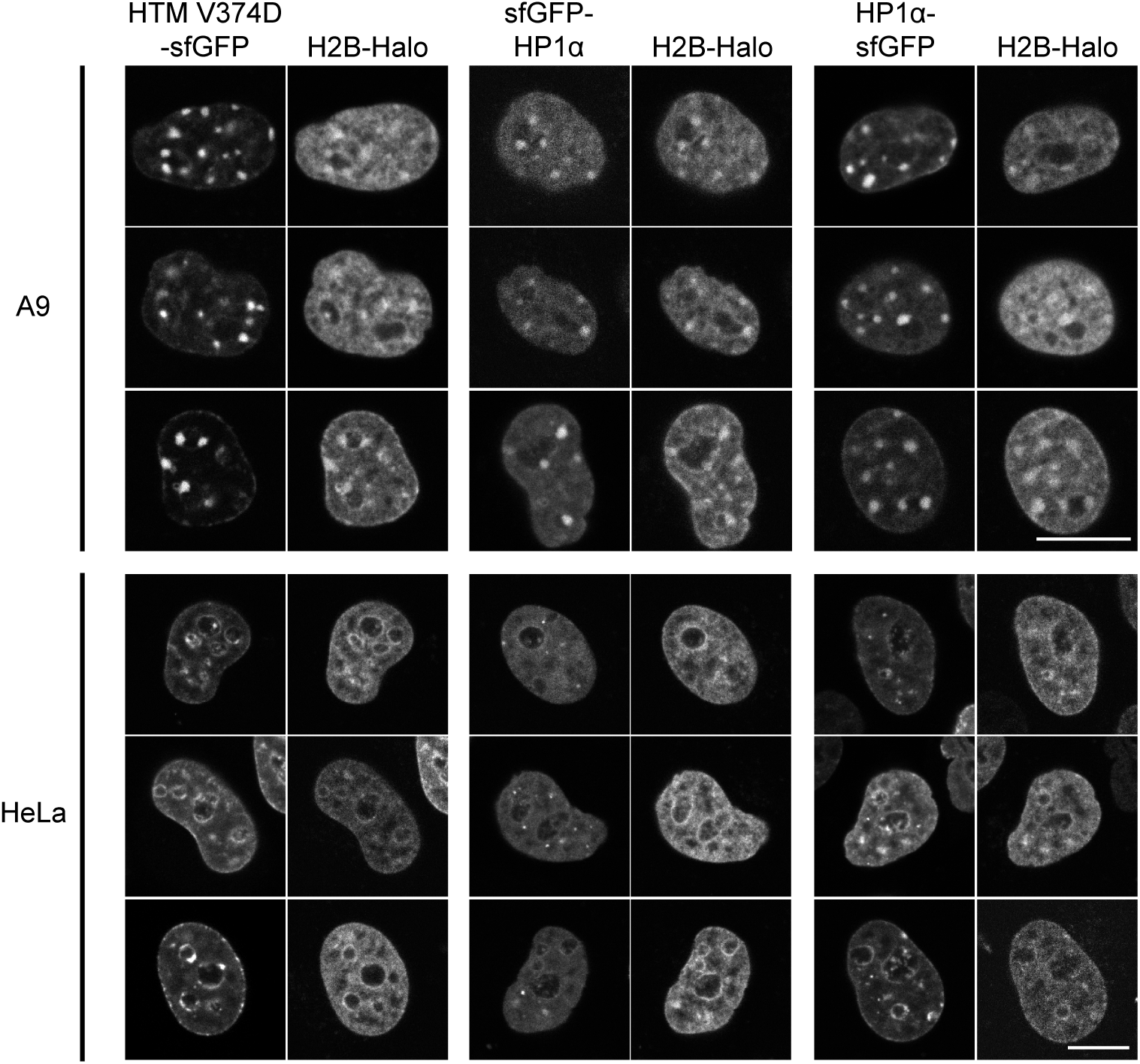
Panels of cells expressing HTM V374D-sfGFP, sfGFP-HP1α, and HP1α-sfGFP. Three nuclei from living A9 and HeLa cells expressing HTM V374D-sfGFP, sfGFP-HP1α, and HP1α-sfGFP, along with H2B-Halo, are displayed. These images provide additional support for the localization patterns of in single nuclei shown in Fig. 6 A and 6B. Scale bars: 10 µm.

**Supplementary Fig. S8.**
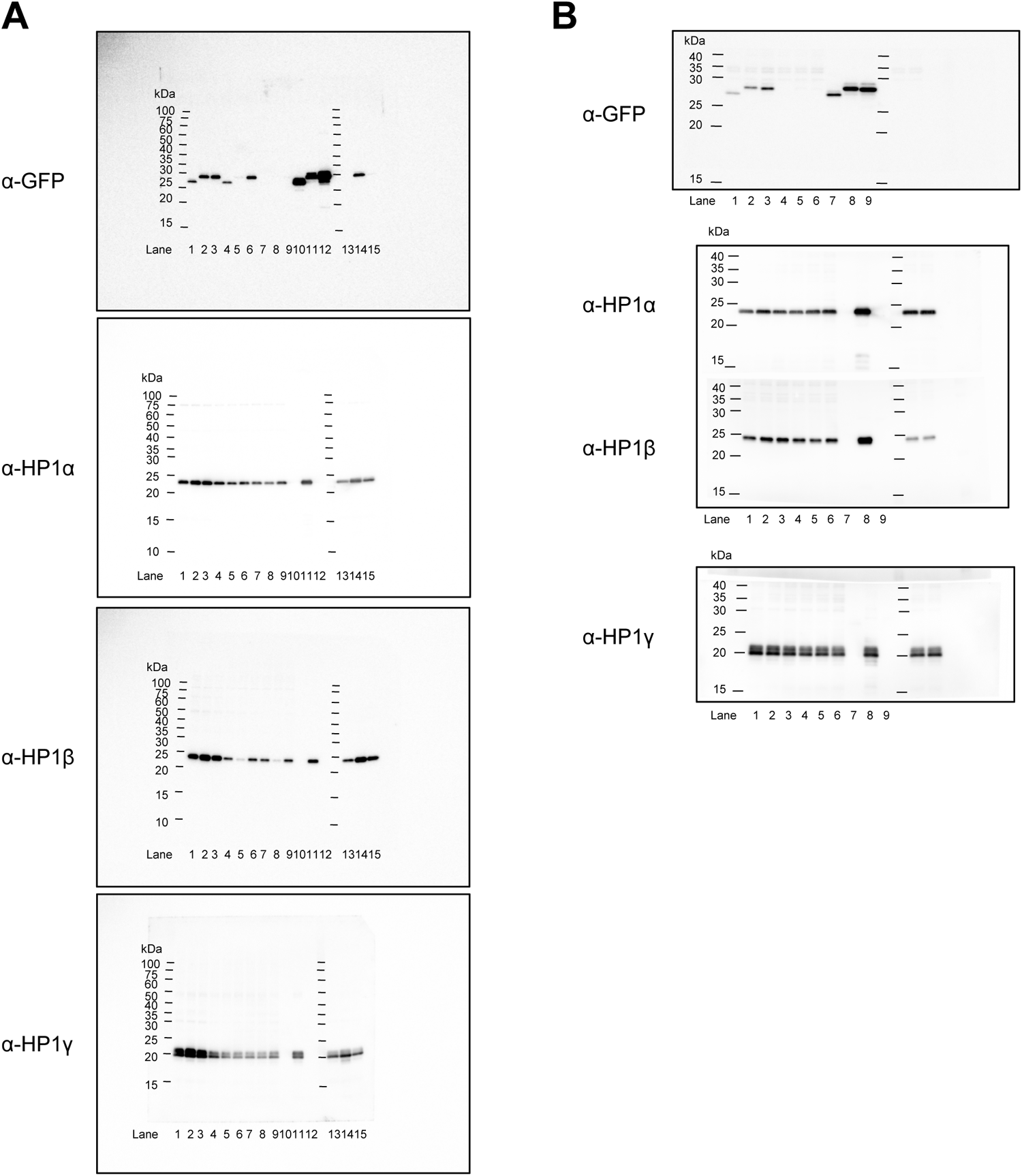
Whole membranes of western blotting. The entire membranes from Western blotting experiments shown in Fig. 2A (A) and 2B (B) are displayed. The positions of the size standards are indicated.

## Legends to Movies

Movie 1. This movie showcases a cell expressing HTM V374D-sfGFP, mCherry-PCNA, and H2B-Halo. It features single confocal images that were acquired at 5 min intervals.

Movie 2. This movie presents enlarged views of a chromocenter in a cell expressing HTM V374D-sfGFP, mCherry-PCNA, and H2B-Halo. It focuses on detailed views of a chromocenter from cells shown in Movie 1.

